# ALS-related p97 R155H mutation disrupts lysophagy in iPSC-derived motor neurons

**DOI:** 10.1101/2023.06.21.545956

**Authors:** Jacob A. Klickstein, Michelle A. Johnson, Pantelis Antonoudiou, Jamie Maguire, Joao A. Paulo, Steve P. Gygi, Chris Weihl, Malavika Raman

**Author notes:** Address correspondence to Malavika Raman.

## Abstract

Mutations in the AAA+ ATPase p97 (also known as valosin containing protein, VCP) cause multisystem proteinopathy 1 (MSP-1) which includes amyotrophic-lateral sclerosis (ALS); however, the pathogenic mechanisms that contribute to motor neuron loss in familial ALS caused by p97 mutations remain obscure. Here, we use two distinct induced pluripotent stem cell models differentiated into spinal motor neurons to investigate how p97 mutations perturb the motor neuron proteome. Using multiplexed quantitative proteomics in these cells, we find that motor neurons harboring the p97 R155H mutation have deficits in lysosome quality control and the selective autophagy of lysosomes (lysophagy). p97 R155H motor neurons are unable to efficiently clear damaged lysosomes and have reduced viability. Additionally, lysosomes in mutant motor neurons have increased pH compared to their wildtype counterparts. The endo-lysosomal damage repair (ELDR) complex is required for clearance of damaged lysosomes and involves UBXD1-p97 interaction which is disrupted in mutant motor neurons. Finally, we report that inhibition of the ATPase activity of p97 R155H using the ATP competitive inhibitor CB-5083 rescues lysophagy defects in mutant motor neurons. These results add to the increasing evidence that endo-lysosomal dysfunction is a key aspect of disease pathogenesis in p97-related disorders.

## Introduction

p97 (also known as valosin containing protein, VCP or CDC48p in yeast) is an evolutionarily conserved type II AAA+ ATPase with important roles in ubiquitin dependent protein quality control. p97 is ubiquitously expressed and forms a homohexamer wherein each monomer is composed of two ATPase domains (D1 and D2) as well as an N-terminal regulatory domain (NTD)^1^. ATP hydrolysis by p97 has been demonstrated to enable substrate unfolding by threading the ubiquitylated substrate through the central pore of the hexamer^2, 3^. This ‘unfoldase’ activity is particularly suited for the extraction of substrates from membranes and multi-protein complexes prior to proteasomal degradation^4^. Key to p97 function and specificity is its ability to form distinct complexes with a host of adaptor proteins. Over 40 adaptors have been identified that enable p97 recruitment to ubiquitylated substrates^5^. We and others have shown that p97-adaptor complexes are important for a large number of cellular processes including organelle contact sites^6^, cell cycle regulation^7^, DNA damage response^8^, and autophagy^9^ (see ^9, 10^ for review). As a result, loss of p97 impacts the degradation of a large cohort of the ubiquitin-modified proteome.

In addition to proteasomal degradation, p97 plays an important role in the endolysosomal system and autophagy^11, 12^ with recent reports highlighting its necessity in maintaining lysosome homeostasis via lysophagy, the autophagic process of turning over damaged lysosomes^13–15^. p97 mediates removal of ubiquitylated intermediates on the lysosomal membrane allowing for autophagy to progress^15^. This action is performed in conjunction with a trimeric adaptor complex composed of the adaptors UBX domain protein 1 (UBXD1), YOD1 deubiquitinase (YOD1), and phospholipase A2 activating protein (PLAA), collectively known as the Endo-Lysosomal Damage Repair (ELDR) complex^13^. Knockdown of p97 or any of the ELDR complex components prevents removal of damaged lysosomes promoting cell death.

Mutations in p97 cause a heterogenous disease known as multisystem proteinopathy 1 (MSP-1) which includes Paget’s disease of the bone, inclusion body myopathy, frontotemporal dementia, and amyotrophic lateral sclerosis (ALS)^16, 17^. Tissue from patients with p97 mutations contain TDP-43 and ubiquitin positive inclusions linking p97-related disease with sporadic forms of these diseases^18^. Certain disease-related p97 mutations have increased ATPase activity in vitro^19^; however, whether this translates to a toxic gain-of-function or a dominant negative effect remains debated^8–10^. The majority of mutations found in humans occur in the NTD and N-terminal-D1 linker region which is important for binding to some classes of adaptors^20^. Indeed, these mutations alter adaptor binding, with some adaptors having increased association (e.g. UFD1-NPL4^21^) and others with decreased association (e.g. UBXD1^22^). Thus, for some p97 mutations, there may be loss of targeting to some substrates leading to their accumulation or precocious degradation of others due to increased targeting; however, this has not been exhaustively tested in relevant cell types.

The cellular processes affected by mutant p97 that lead to disease are still intensely debated. Models in immortalized cell types such as U2OS and NSC-34 have found defects in lysophagy leading to an accumulation of damaged lysosomes and increased cell death^13, 23^. Drosophila and rodent models of p97 disease have shown defects in protein degradation, autophagy, and mitochondrial function^24–26^. Induced pluripotent stem cell (iPSC) - derived motor neuron models of p97 disease have heterogenous phenotypes as well. Initial studies utilizing iPSC-derived motor neurons from a patient with the R155H mutation found increased levels of TDP-43, ubiquitin, and autophagy-related proteins in whole cell lysate^27^. Patani and colleagues refined this phenotype finding TDP-43 mislocalization, ER stress induction, and mitochondrial dysfunction in motor neurons from patients carrying the R155C and R191Q mutations^28^. Differences in genetic backgrounds between patient and control iPSCs can make direct comparisons challenging. However, a recent study overcame this issue by introducing the R155H mutation into wildtype iPSCs and correcting the R155H mutation in patient derived iPSCs to the wildtype allele to create isogenic controls. Interestingly, this approach found no changes in ER stress or TDP-43 mislocalization; however, aberrant cell-cycle regulation was determined to be a contributor to decreased viability^29^. A common feature seen in both cellular and animal models is decreased viability in p97 mutant neurons. Thus, further studies are needed to elucidate features that may be common between different models. Additionally, to the best of our knowledge there has been no study of lysophagy nor the effect of p97 mutation on lysophagy in human motor neurons.

To begin to understand how p97 mutations impact the cellular proteome, we performed quantitative proteomics on two distinct p97 R155H iPSC-derived motor neuron cell lines. The first iPSC line was derived from a patient harboring the R155H mutation that was then corrected to the wildtype allele using CRISPR/cas9 editing. The second iPSC line was created as part of the iPSC Neurodegenerative Disease Initiative (iNDI), wherein the R155H mutation was introduced into the endogenous p97 genomic locus of KOLF2.1 cells to create both heterozygous and homozygous lines^30, 31^. These two cell-based models allow us to interrogate how the cellular proteome is impacted by p97 mutation alone (KOLF2.1 lines) as well as how patient specific genetic backgrounds modify the p97 R155H phenotype. From our proteomic studies, we found that each genetic background had over 200 differentially expressed proteins, but only a few proteins were common between the patient and KOLF2.1 cell lines. For example, KOLF2.1 R155H cells had a pronounced defect in mitochondrial homeostasis; surprisingly, the patient-derived cells did not have this phenotype. An unbiased clustering analysis identified autophagy and lysosomal proteins as the most significantly altered in both the mutant lines. Of the few commonly altered proteins, we identified multiple components of lysosome repair machinery, namely ANXA1, ANXA2, and TRIM16.

To investigate a potential lysophagy defect in our models, we utilized the lysosomal damaging agent L-leucyl leucine-O-methyl ester (LLOME). Following LLOME treatment, we found that while wildtype motor neurons were able to effectively clear damaged lysosomes, p97 mutant neurons were not. In wildtype and mutant neurons LLOME treatment resulted in robust ubiquitylation of lysosomes that recruited both wildtype and mutant p97. Interestingly, while LLOME treatment stimulated the association of wildtype p97 with UBXD1, this complex was less effectively formed by p97 R155H suggesting ineffective ELDR complex formation. Because mutant p97 has increased ATPase activity, previous studies have utilized p97 inhibitors to reverse disease phenotypes^29, 32, 33^. We co-treated motor neurons with LLOME and CB-5083, a highly specific competitive p97 inhibitor, and found that inhibition of mutant p97 rescued lysophagy defects and prevented LLOME-induced cell death. Overall, our studies suggest that p97 R155H mutation negatively impacts lysosome quality control in human motor neurons.

## Results

### Efficient differentiation of two isogenic iPSC R155H p97 models into functional motor neurons

We used two orthogonal models to investigate how the p97 R155H mutation impacted the motor neuron proteome. The first was a wildtype KOLF2.1 iPSC line that was CRISPR edited to introduce the R155H mutation into the endogenous p97 locus that was developed by iNDI^30^ (Figure 1A). We used two clonal lines of both heterozygous and homozygous mutants for our studies (referred to as KOLF2.1 wildtype (WT), heterozygous (Het), and homozygous (Hom)). The second was the 392.1 iPSC line from a forty eight year old male diagnosed with ALS, dementia and Paget’s disease, harboring the R155H p97 mutation. The R155H mutation was reverted to wildtype by CRISPR/CAS9 editing (referred to as 392.1 wildtype (WT) and R155H (Figure 1A). Genotypes for both cell lines were confirmed by DNA sequencing (Figures S1A and S2A). A normal karyotype was confirmed for both parental lines (Figures S1B and S2B). Wildtype and mutant iPSCs were indistinguishable from each other in colony morphology, the ability to form embryoid bodies, and the expression of pluripotency markers (Figures 1B, S1C, S2C, and S2D). We established a spinal motor neuron differentiation protocol using previous studies^34–36^. Briefly, iPSCs were induced to neural ectoderm fate using dual-SMAD inhibition and a WNT pathway agonist (Figure 1C). After three days, retinoic acid (RA) and smoothened agonist (SAG, an activator of the SHH pathway) were added to create NES+, SOX2+, PAX6+ neural precursor cells (NPCs) (Figures 1D and E, S1D, S2E, and S2F). At this stage, these cells can be expanded and banked. NPCs were further patterned using RA and SAG to a ventral, caudal fate for three days to create motor neuron progenitors (Figure 1C). Progenitors were replated onto poly L-ornithine (PLO), laminin, and fibronectin coated plates and cultured in motor neuron maturation media containing neurotrophic factors (neurotrophin-3 (NT-3), brain derived neurotrophic factor (BDNF), and glial derived neurotrophic factor (GDNF)), RA, SAG, and the gamma-secretase inhibitor, compound E to activate the notch pathway. 21 days of maturation produced >90% class III ý-tubulin (TUJ1), neurofilament heavy chain (SMI32), choline acetyltransferase (ChAT) positive cells with elongated processes typical of motor neurons (Figure 1D and E, S1E, S2G).

**Figure 1.**
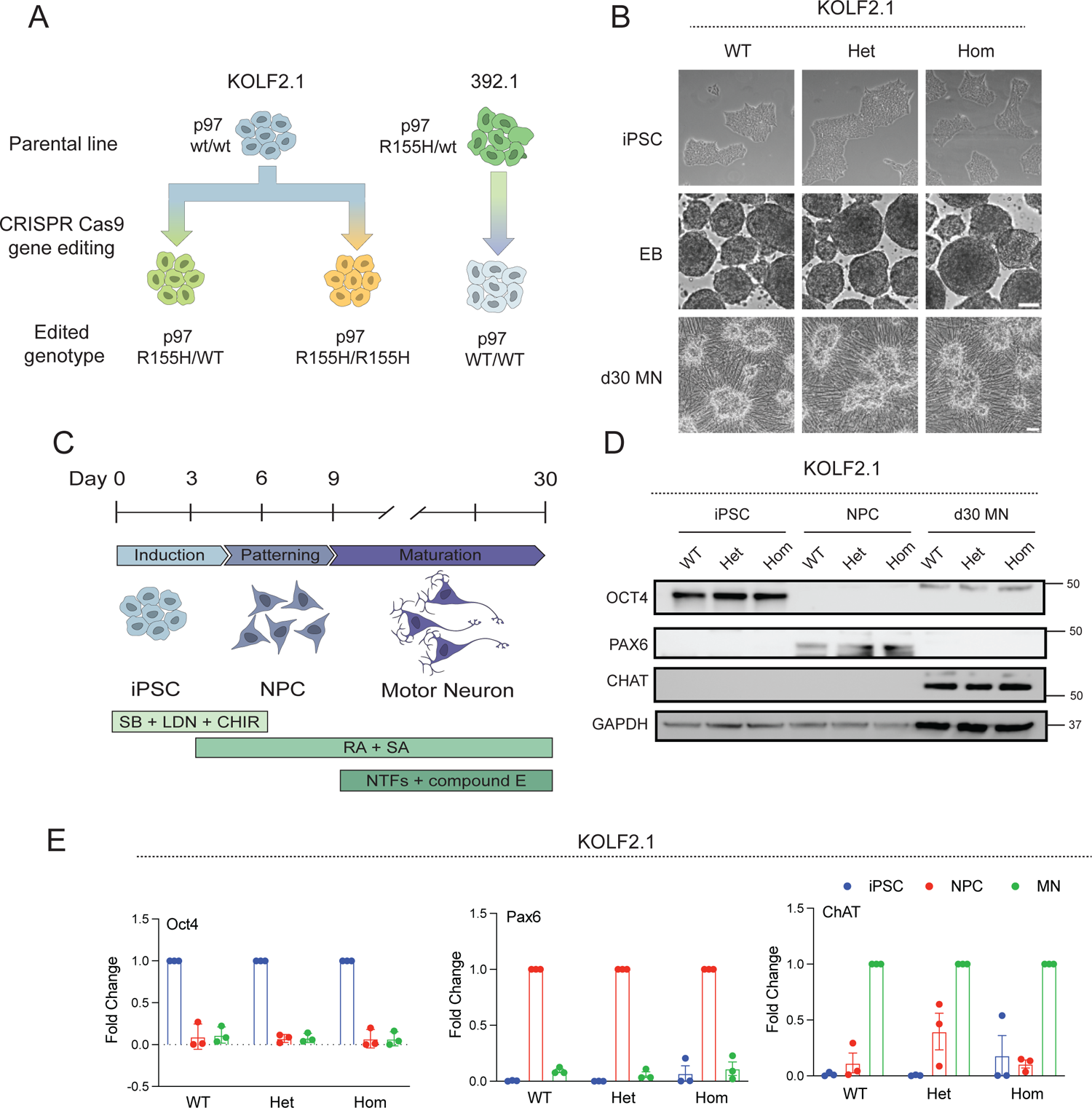
Characterization of IPSC-derived motor neurons. A. Two isogenic iPSC models were generated using gene editing to introduce the R155H p97 in the endogenous locus (KOLF2.1) and correct the R155H mutation (392.1). B. Phase contrast images of iPSCs (top row), embryoid bodies (middle row), and day 30 motor neurons (bottom row). Scale bars 20µm (top and bottom rows) and 100µm (middle row). C. Schematic of the motor neuron differentiation protocol. D. Immunoblot of iPSCs, NPCs, and day 30 motor neurons, and their respective cell markers demonstrate efficient differentiation of wildtype and mutant p97 cells. E. qPCR of WT, heterozygous, and homozygous KOLF2.1 iPSCs, NPCs, and motor neurons and markers for each stage. EB – embryoid body, MN – motor neuron, NPC – neural precursor cell, SB - SB431542, LDN - LDN193189, CHIR - CHIR99021, RA – retinoic acid, SA – smoothened agonist, NTFs – neurotrophic factors.

### Characterization of cellular phenotypes in p97 R155H motor neurons

We next sought to characterize several cellular phenotypes attributed to mutant p97 in these cell lines. We confirmed that mature KOLF2.1 motor neurons were functional using Fluo-4 AM live-cell calcium imaging which showed >90% of cells responded to 80mM KCl depolarization (Figure 2A and B). Interestingly, KOLF2.1 heterozygous and homozygous motor neurons had lower maximal calcium transients in response to KCl suggesting altered calcium dynamics (Figure 2B, right). We next performed whole-cell patch clamp in wildtype and homozygous KOLF2.1 motor neurons at day 20. Both genotypes had equivalent input resistance and had both spontaneous action potentials (APs) and trains of action potentials in response to depolarizing current (Figure 2C, left and middle); however, homozygous motor neurons trended towards increased AP frequency though this did not reach statistical significance (Figure 2C right). These data suggest that our differentiation process produces functional motor neurons and that p97 mutation impacts their electrical activity as reported by other groups^28, 29^.

**Figure 2.**
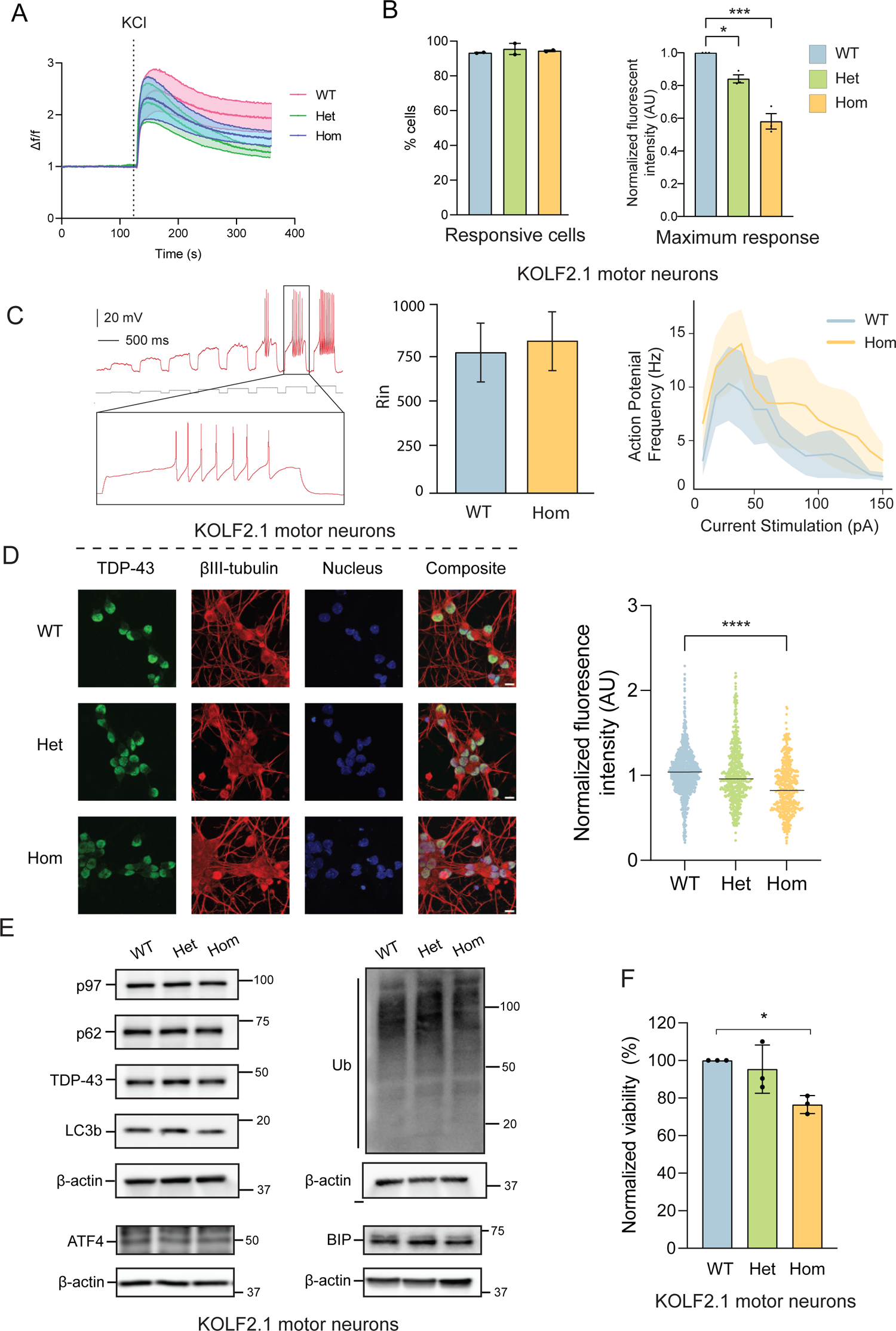
R155H p97 motor neurons recapitulate ALS phenotypes. A. Representative average Fluo-4 fluorescence values from KOLF2.1 motor neurons after 80 mM KCl stimulation (dotted line). The shaded area represents the 95% confidence interval. N = >50 cells per condition. B. Percent of KOLF2.1 motor neurons responsive to KCl stimulation (left). Maximal normalized Fluo-4 fluorescence in motor neurons. N = >150 cells over 3 independent experiments per condition. Data are expressed as mean ± SEM. *p<0.05, ***p<0.001; One-way ANOVA with Dunnett’s multiple comparison test. C. A sample trace from a WT KOLF2.1 motor neuron in response to a series of depolarizing current steps via whole cell patch clamp (left). Input resistance from WT and homozygous motor neurons (middle). Input-output curves of motor neurons in WT and homozygous motor neurons at day 20 (right). D. Representative images of TDP-43 immunofluorescence in KOLF2.1 motor neurons (left panels). Quantification of nuclear TDP-43 fluorescence (right). N = >5000 cells over 4 independent experiments. Data points represent individual cells; lines represent means. ****p<0.0001; one-way ANOVA with Dunnett’s multiple comparison test. Scale bar 10µm. E. Representative immunoblots of TDP-43, autophagy markers (p62, LC3b), ER stress markers (ATF4, BIP), and total ubiquitylated proteins (Ub) in KOLF2.1 motor neurons. N = 3 independent experiments. F. Normalized viability of KOLF2.1 motor neurons at day 30. Data expressed as means ± SEM. *p<0.05; one-way ANOVA with Dunnett’s multiple comparison test.

TAR DNA binding protein 43 (TDP-43) is a nuclear RNA binding protein that regulates the splicing of hundreds of genes^37^ and its mislocalization to the cytosol and subsequent aggregation is prominent feature in ALS and MSP-1^18^. We investigated TDP-43 localization in our iPSC lines using confocal microscopy and found that both mutant 392.1 and KOLF2.1 motor neurons had depleted levels of TDP-43 in the nucleus compared to their wildtype counterparts (Figure 2D and S3A). However, total TDP-43 levels were equivalent between lines suggesting that nuclear depletion was not due to wholesale TDP-43 depletion (Figures 2E and S3B).

A key cellular function of p97 is ER associated degradation (ERAD) which is the retrotranslocation of misfolded, ubiquitylated proteins from the ER for proteasomal degradation. Whether p97 mutation negatively impact ERAD is debated as several studies have linked p97 mutations to induction of ER stress pathways^28^ whereas others found no change in mutant iPSC-derived motor neurons^29^. In our models, we found no change in several markers of ER stress including the transcription factor ATF4 or the ER chaperone BIP (Figure 2E and S3B). Furthermore, we found no significant change in total ubiquitin conjugates or autophagy markers p62 or LC3b (Figure 2E and S3B).

Finally, we measured cell viability and found that KOLF2.1 p97 R155H homozygous motor neurons had decreased survival relative to wildtype. The 392.1 patient line also had a significant though modest decrease in cell viability in agreement with previous studies in similar models^28, 29^ (Figure 2F and S3C).

### Unbiased quantitative proteomics reveals distinct proteome alterations between KOLF2.1 and patient cell lines

To interrogate changes in motor neurons due to p97 mutations at the proteome level, we employed unbiased quantitative proteomics using tandem mass tags (TMT)^38^. Wildtype and R155H motor neurons were aged 30 days from either KOLF2.1 or 392.1 and then harvested, digested, and peptides were labelled with tandem mass tags and analyzed by liquid chromatography and mass spectrometry (LC/MS-MS) (Figure 3A, Supplementary Table 1)^39, 40^. More than 200 proteins were significantly altered (Log_2_ fold change mutant:WT > 0.7) between mutants and wildtype in both cell lines suggesting broad changes at the protein level despite relatively mild phenotypic alterations (Figure 3B-D). Gene ontology (GO) analysis of differentially expressed proteins (DEPs) found different pathways altered in KOLF2.1 and 392.1 mutant motor neurons despite harboring the same R155H mutation (Figure S4A-C). Notably, KOLF2.1 homozygous motor neurons showed significant depletion of mitochondrial proteins (Figure S4B). In particular, depletion of proteins that are part of complexes regulating import of proteins into mitochondria, and subunits of electron transport chain (ETC) complexes I-IV was observed (Figure S4D). These results were further validated by immunoblot for components of the ETC (TMEM70 and MT-ND2) and the translocase complex (TOMM20 and TOMM70) (Figure S4E). In agreement with significant perturbation to the mito-proteome, KOLF2.1 mutant motor neurons had significantly decreased mitochondrial membrane potential as measured using MitoTracker red (Figures S4F and G). No significant changes were observed in mitochondrial morphology (Figure S4H). Interestingly, these mitochondrial changes were not seen in the 392.1 mutant motor neurons (Figure S4C, and data not shown). Therefore, we sought to identify altered pathways that were shared between KOLF2.1 and 392.1 motor neurons. We utilized weighted gene correlation network analysis (WGCNA) adapted for proteomic studies to segment proteins into modules with similar expression profiles in an unbiased manner^41^. This methodology uses a dynamic branch cutting algorithm to separate clusters identified via a topological overlap matrix into distinct modules (Figure S5A, Supplementary Table 2). Inspection of module eigenprotein expression revealed agreement between KOLF2.1 and 392.1 wildtype lines, but a divergence of expression pattern in the mutant neurons (Figure S5B). To identify modules that represented proteins changing in a similar manner between cell lines, modules were ranked by a composite score based on concordance between genetic backgrounds and average log_2_ fold change (Figure S5 D-F, Supplementary Table 2). The member proteins of the top three ranked modules were used for GO analysis to identify enriched terms. Multiple GO terms related to autophagy and lysosomal homeostasis were enriched in the top modules (Figure S5D and E middle panels).

**Figure 3.**
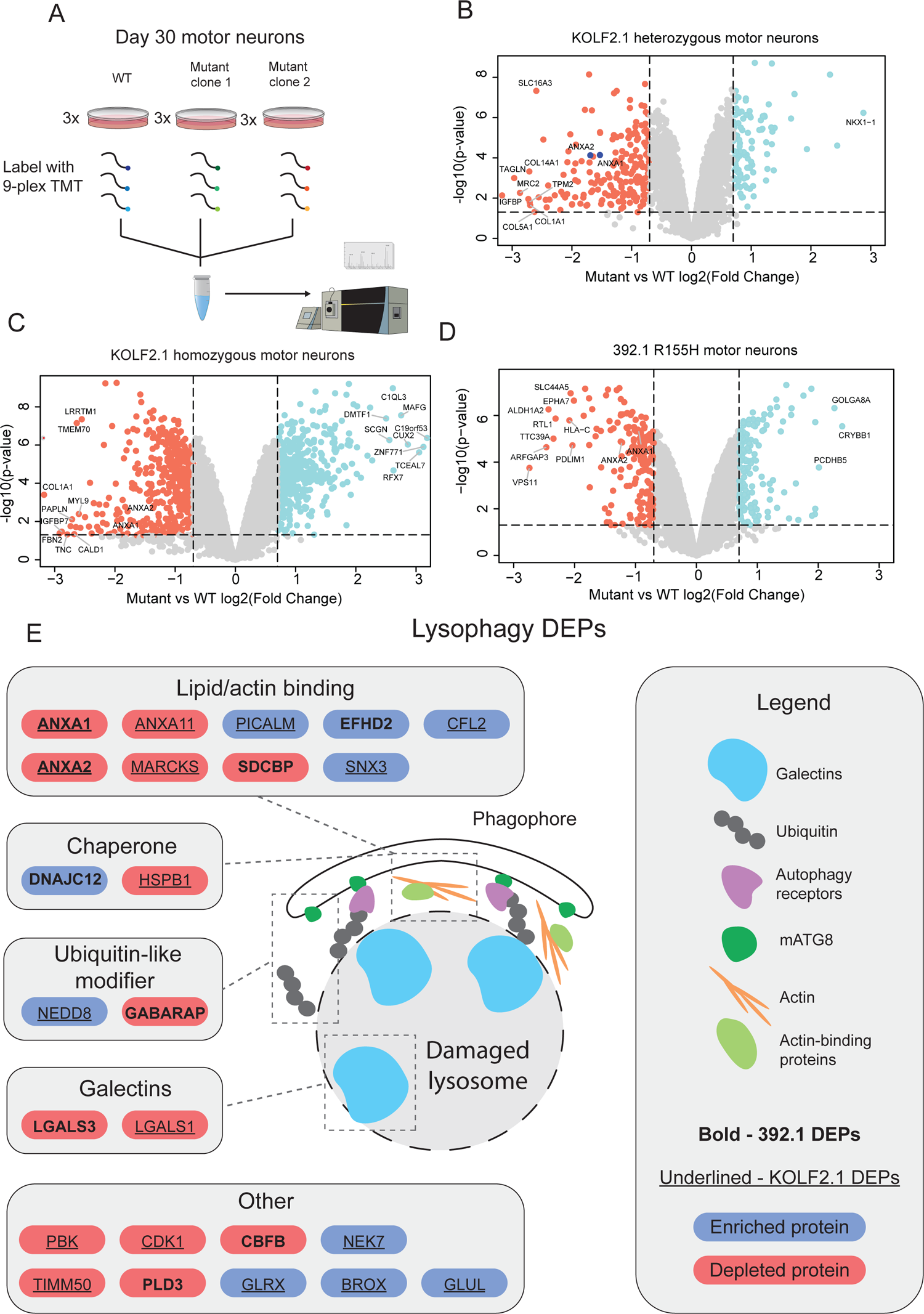
Quantitative proteomics identifies autophagy defects in R155H motor neurons. A. Schematic of the experimental setup for the KOLF2.1 cell lines. Three replicates each of WT and 2 clones of heterozygous or homozygous R155H p97 motor neurons were used in two 9-plex TMT experiments. Three replicates of WT and R155H p97 392.1 motor neurons were used for a 6-plex TMT experiment. B-D. Volcano plots for the indicated cell line. Cutoff values of 0.7 log_2_ fold change and 0.05 p-value were used to determine differentially expressed proteins (DEPs). E. Lysophagy related proteins that were significantly altered in R155H motor neurons. Red and blue backgrounds indicate depleted and enriched proteins respectively.

The identification of several pathways relating to lysosomal homeostasis was particularly interesting as p97 plays a critical role in the turnover of lysosomes and depletion of p97 leads to the persistence of damaged lysosomes^13^. We compared our proteomics data set to two published studies that identified proteins enriched on damaged lysosomes^42^ or were ubiquityled following lysosomal damage^15^. From this list of lysophagy proteins, we found over 20 DEPs that were differentially modulated in KOLF2.1 and/or 392.1 motor neurons suggesting that p97 R155H hinders the ability of motor neurons to properly respond to damaged lysosomes (Figure 3E).

### Damaged lysosomes in motor neurons preferentially recruit LGALS8

The alterations in autophagy and endocytosis pathways prompted us to investigate lysosomal turnover in p97 mutant motor neurons. L-leucine leucyl-O-methyl ester (LLOME) is a lysomotropic agent that permeabilizes lysosomal membranes. LLOME treatment has been shown to initiate galectin 3 (LGALS3) and autophagy component (p62 and LC3B) recruitment for lysosomal turnover in immortalized cells^43^ and cortical neurons^42^. Galectin recruitment to lysosomes is recognized as a sensitive measure of lysosomal damage^44^; however, to our knowledge, lysophagy and markers of damaged lysosomes have not been evaluated in human spinal motor neurons. In addition to LGALS3, other galectins such as LGALS1, LGALS8, and LGALS9 have also been shown to associate with lysosomes following LLOME treatment^45^. We evaluated the recruitment of LGALS 1, 3, 8 and 9 following LLOME treatment in both NPCs and d30 motor neurons. Surprisingly, we found that while NPCs recruited both LGALS3 and LGALS8 to damaged lysosomes (Figure 4A), motor neurons selectively recruited LGALS8 (Figure 4B). This is in agreement with single-cell expression data of spinal cord cell types^46^ that indicated that LGALS8 has highest expression in motor neurons compared to other galectins (Figure 4C). We note that in addition to punctate localization, LGALS8 immunostaining also showed a non-punctate linear pattern both in untreated and treated conditions (Figure 4B). To ensure that LGALS8 puncta formed in damaged lysosomes, we performed Airyscan super resolution microscopy on motor neurons stained for the lysosomal protein LAMP1 and LGALS8 after LLOME treatment. All LGALS8 puncta colocalized with LAMP1 while the linear ‘dashes’ did not (Figure 4B right panels). LGALS8 puncta also colocalized with ubiquitin demonstrating that LGALS8 faithfully represents damaged lysosomes (Figure 4D). Non-punctate LGALS8 structures were not quantified in our subsequent studies.

**Figure 4.**
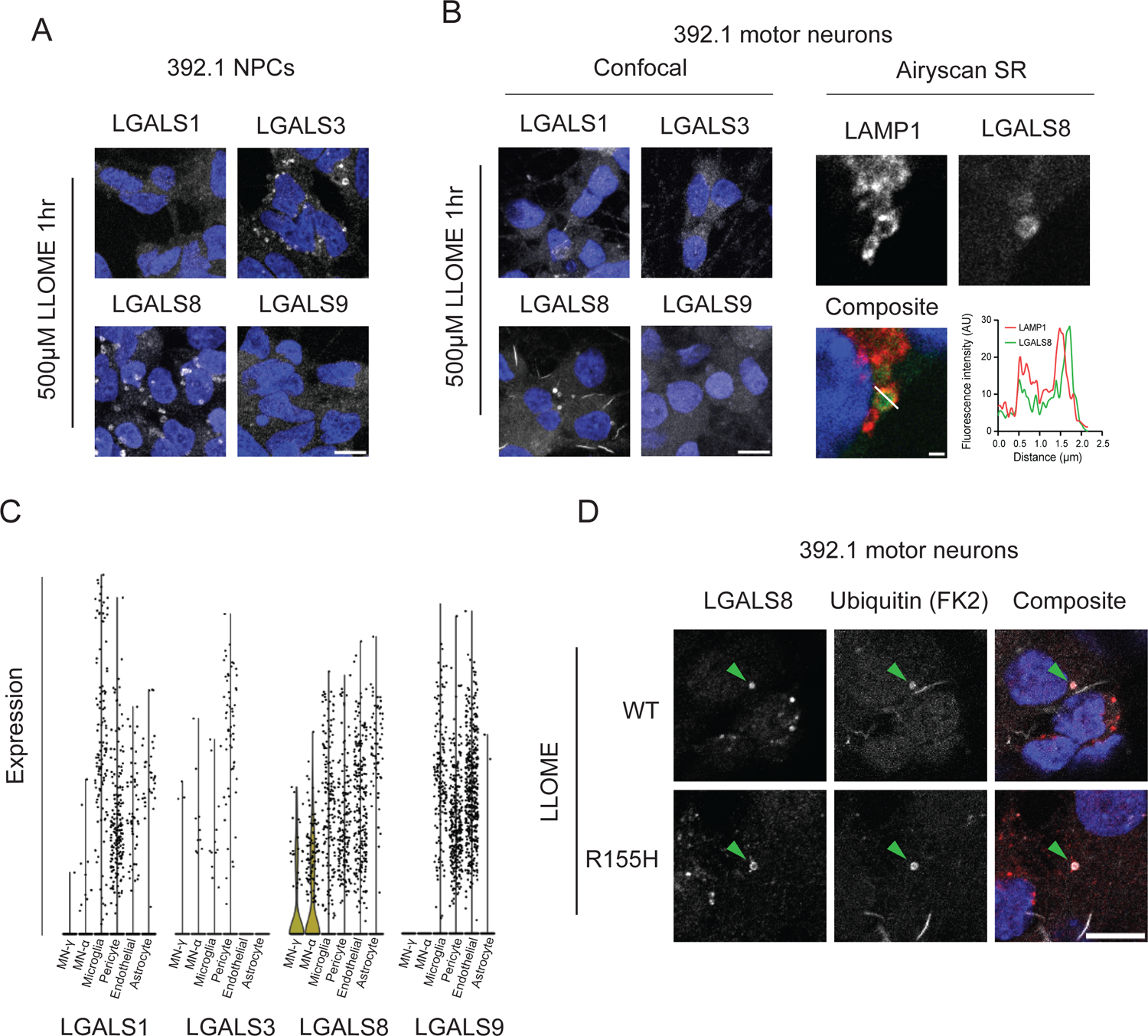
Motor neurons preferentially recruit LGALS8 to ubiquitylated lysosomes. A. Representative images of LGALS1, LGALS3, LGALS8, and LGALS9 immunofluorescence in neural precursor cells (NPCs) following 1hr of 500µM LLOME treatment. N = 2 independent experiments. Scale bar 10µm. B. Representative images of LGALS1, LGALS3, LGALS8, and LGALS9 immunofluorescence in motor neurons following 1hr of 500µM LLOME (left). Airyscan images of motor neurons following LLOME treatment co-stained with LAMP1 and LGALS8 show colocalization (right). N = 2 independent experiments. Scale bars 10µm (left) and 1µm (right). C. Single-cell RNAseq expression data of LGALS1, LGALS3, LGALS8, and LGALS9 in spinal cord tissue from the SeqSeek dataset^46^. Data are expressed as violin plots of normalized transcripts per million (TPM) values. Individual data points represent outliers. D. Representative images of LLOME treated motor neurons co-stained for LGALS8 and ubiquitin demonstrating colocalization. N = 3 independent experiments. Scale bar 10µm. LLOME – L-leucyl leucine O-methyl ester.

### R155H p97 disrupts lysophagy in motor neurons

As LGALS8 was the most robust marker for lysosomal damage in motor neurons, we utilized LGALS8 puncta formation as a measure of lysosomal damage in the following studies. We next asked whether motor neurons harboring R155H p97 had defects in lysophagy. Motor neurons were treated with LLOME and then allowed to recover for 5 or 24 hours and lysosome repair was monitored by LGALS8 recruitment (Figure 5A). Wildtype and mutant motor neurons had an equal number of LGALS8 puncta following treatment; however, after 5 hours of recovery, wildtype neurons had completely cleared damaged lysosomes while the mutants had LGALS8 puncta that persisted 24 hours after treatment (Figures 5B, 5D left, and 5E left). p62/SQSTM1 is a known autophagy receptor that is recruited to damaged lysosomes via interaction with ubiquitylated cargo^47^. Immunofluorescence for p62 demonstrated equal recruitment 5 hours after treatment between wildtype and mutant; however, mutant motor neurons had significantly increased p62 puncta 24 hours after treatment compared to wildtype (Figures 5C, 5D right, and 5E right). Failure to clear damaged lysosomes can lead to cell death^48^. We assessed viability following LLOME treatment and found that mutant motor neurons had increased cell death following release while wildtype neurons had no change (Figures 6A and S6A). We next asked whether disrupted levels of proteins involved in lysosome repair may also impact lysosome function at basal levels even in the absence of overt lysosomal damage. We therefore interrogated basal lysosomal function using the pH sensitive lysosomal dye LysoSensor DND-189 which accumulates in acidic compartments fluoresces at low pH. To confirm the sensitivity of this dye, we performed live cell imaging on motor neurons after 30 minutes of bafilomycin-A1, a known inhibitor of lysosomal vacuolar H+ ATPase or LLOME. Following subthreshold LLOME or bafilomycin-A1 treatment, we found that the LysoSensor fluorescence was significantly decreased demonstrating compromised pH (Figures 6B and S6B). Notably, mutant motor neurons had decreased fluorescence at basal levels compared to wildtype (Figure 6B and S6B).

**Figure 5.**
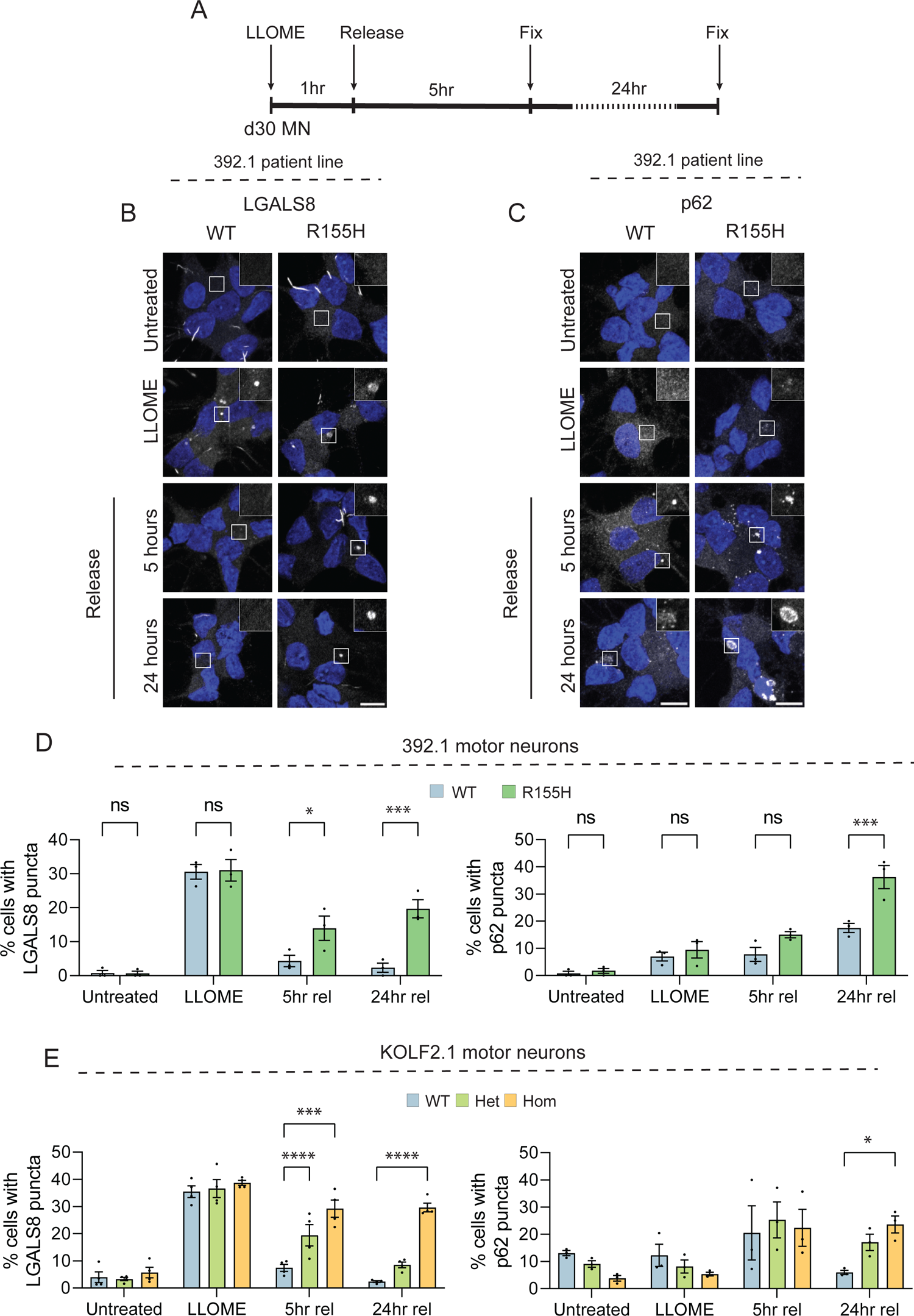
R155H p97 motor neurons are unable to clear damaged lysosomes and accumulate p62 following LLOME treatment. A. Schematic of experimental design. Day 30 motor neurons were fixed either before LLOME treatment, directly after treatment or 5 and 24 hours after release. B. Representative images of 392.1 motor neurons stained for LGALS8 before, during, and after LLOME treatment. Scale bar 10µm. C. Representative images of 392.1 motor neurons stained for p62. Scale bar 10µm. D. Quantification of B (left) and C (right) in 392.1 motor neurons. Data expressed as means ± SEM. ns – nonsignificant, *p<0.05, ***p<0.001; two-way ANOVA with Sidak’s multiple comparison test. E. Quantification of LGALS8 puncta (left) and p62 puncta (right) in KOLF2.1 motor neurons. Data expressed as means ± SEM. ns – nonsignificant, *p<0.05, ***p<0.001, ****p<0.0001; two-way ANOVA with Dunnett’s multiple comparison test. D30 MN – day 30 motor neurons, LLOME – L-leucyl leucine O-methyl ester. 5hr rel – 5-hour release, 24hr rel – 24-hour release.

**Figure 6.**
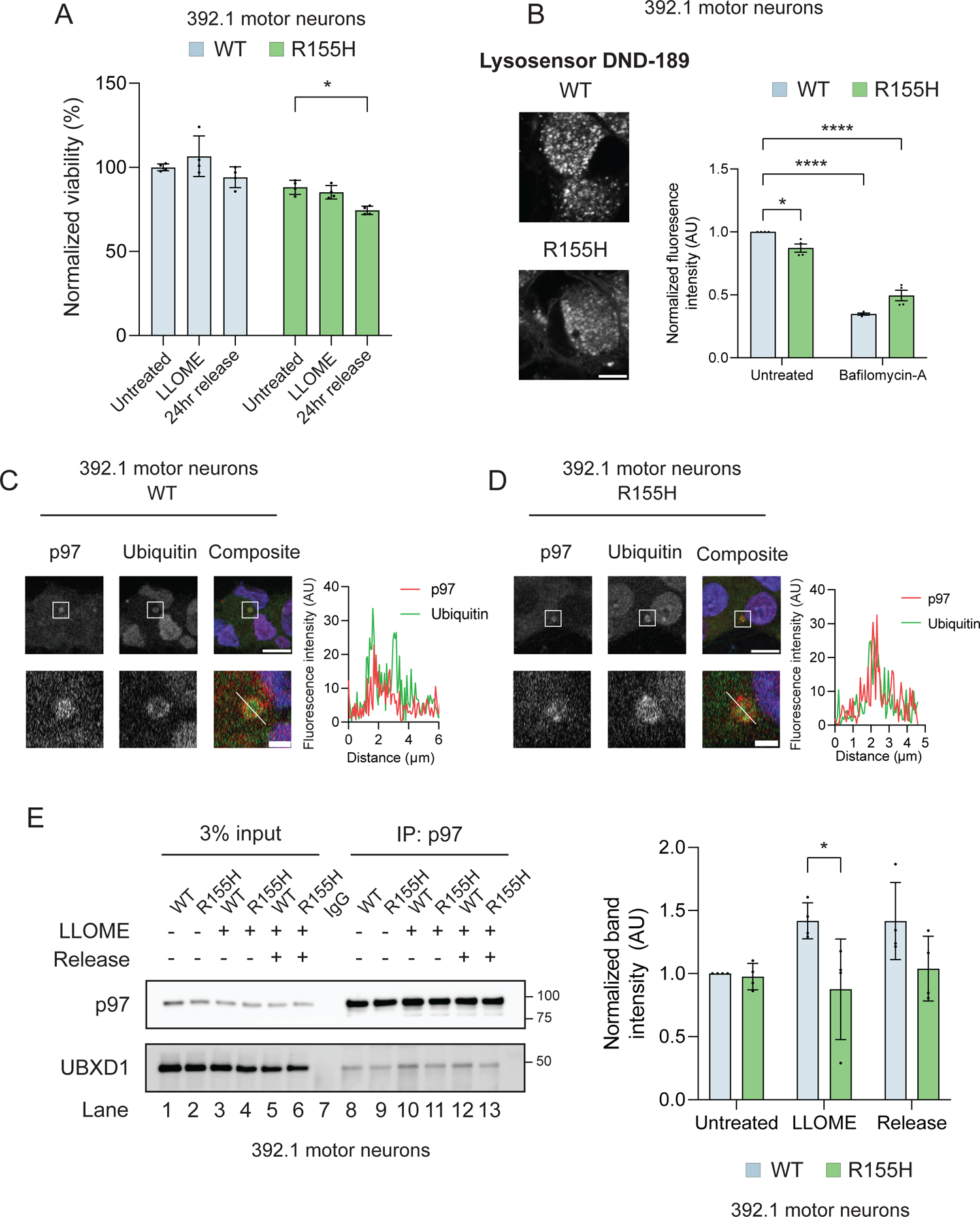
R155H p97 sensitizes motor neurons to lysosomal damage, decreases lysosomal acidity, and disrupts UBXD1 recruitment. A. Normalized viability in 392.1 motor neurons in untreated, LLOME treated or after release. N = 4 independent experiments. Data expressed as means ± SEM. *p<0.01; two-way ANOVA with Dunnett’s multiple comparison test. B. Representative images of Lysosensor DND-189 live cell imaging in 392.1 motor neurons (left). Quantification of DND-189 fluorescence in untreated and bafilomycin-A treated cells. N = 4 independent experiments. Data expressed as means ± SEM. *p<0.05, ****p<0.001; two-way ANOVA with Sidak’s multiple comparison test. Scale bar 5 µm. C-D. Representative images of wildtype (C) and R155H p97 (D) 392.1 motor neurons co-stained with p97 and ubiquitin showing colocalization (left). Line graph of the fluorescence intensity of p97 and ubiquitin as indicated in the bottom right image panel. N = 3 independent experiments. Scale bars 10µm (upper panels) and 1µm (lower panels). E. Immunoblot of endogenous p97 immunoprecipitation in 392.1 motor neurons before, during, and after LLOME treatment (left). Release condition represents 5 hours of recovery. Quantification of UBXD1 band intensities normalized to immunoprecipitated p97 (left). N = 4 independent experiments. Data expressed as means ± SEM. *p<0.05; two-way ANOVA with Sidak’s multiple comparison test.

Ubiquitylation of lysosomal proteins is a key step in lysophagy and is required for lysosomal clearance^49^. To determine if damaged lysosomes in mutant neurons were ubiquitylated, we co-stained for LGALS8 and ubiquitin using the FK2 antibody that recognizes mono- and poly-ubiquitin chains. We found that LGALS8 puncta colocalized with ubiquitin in both wildtype and mutant motor neurons (Figures S6C and D). Knockdown of p97 results in the stalled clearance of damaged lysosomes thus mutations in p97 may prevent the recruitment to damaged lysosomes^13^. However, we found that both wildtype and mutant p97 were equally recruited to ubiquitylated lysosomes (Figures 6C and D). Mutations in p97 have been reported to alter the association with the adaptor UBXD1, a component of the ELDR complex^22^. To test the association of p97 with UBXD1, we immunoprecipitated endogenous p97 from wildtype and mutant motor neurons before, during, and after LLOME treatment. We found that at basal levels, both wildtype and mutant p97 associated with UBXD1; however, during LLOME treatment, wildtype p97 robustly increased association with UBXD1 while mutant p97 did not (Figures 6E, compare lanes 10 to 11 and 12 to 13 and S6E, compare lane 13 to 14 and 15). Taken together, our studies suggest that p97 R155H motor neurons have perturbed lysosomal function and are unable to repair damaged lysosomes upon lysosome membrane permeabilization.

### Inhibition of p97 rescues lysophagy defects in mutant motor neurons and disrupts lysophagy in wildtype neurons

p97 R155H has increased ATPase activity in vitro, and multiple studies have used p97 inhibitors to rescue distinct disease phenotypes including in iPSC-derived motor neurons^29, 32^. However, it is unknown if p97 inhibition rescues defective lysophagy. We attempted to prevent the persistence of damaged lysosomes in mutant motor neurons using CB-5083, a highly specific p97 ATP-competitive inhibitor^50^ (Figure 7A). p97 inhibition by itself did not cause LGALS8 or p62 puncta formation in CB-5083 treated cells (Figures 7B and S7A-E). Notably, CB-5083 treatment caused mutant neurons to accelerate the clearance of damaged lysosomes (monitored by persistence of LGALS8 puncta) in a manner that was comparable to wildtype neurons (Figures 7B, C and S7A, B). Indeed, 24 hours following LLOME treatment, mutant neurons were indistinguishable from wildtype (Figures 7C). While wildtype neurons treated with CB-5083 were able to clear LGALS8 puncta at a similar rate to untreated neurons, they showed persistent and elevated levels of p62 puncta 24 hours after LLOME (Figures S7C-F). Indeed, increased p62 puncta was observed upon LLOME-CB-5053 co-treatment compared to LLOME alone (Figure S7F). This may be a consequence of general perturbations in autophagy due to p97 inhibition. We next asked if p97 inhibition could rescue lysosomal pH in mutant motor neurons. CB-5083 treatment restored basal lysosomal pH in mutant motor neurons to that of wildtype neurons (Figures 7D, E and S7G). Furthermore, CB-5083 treatment prevented cell death in mutant motor neurons during release from LLOME (Figures 7F and S7H). Thus, we conclude that p97 inhibition partially rescues defective lysophagy due to mutant p97.

**Figure 7.**
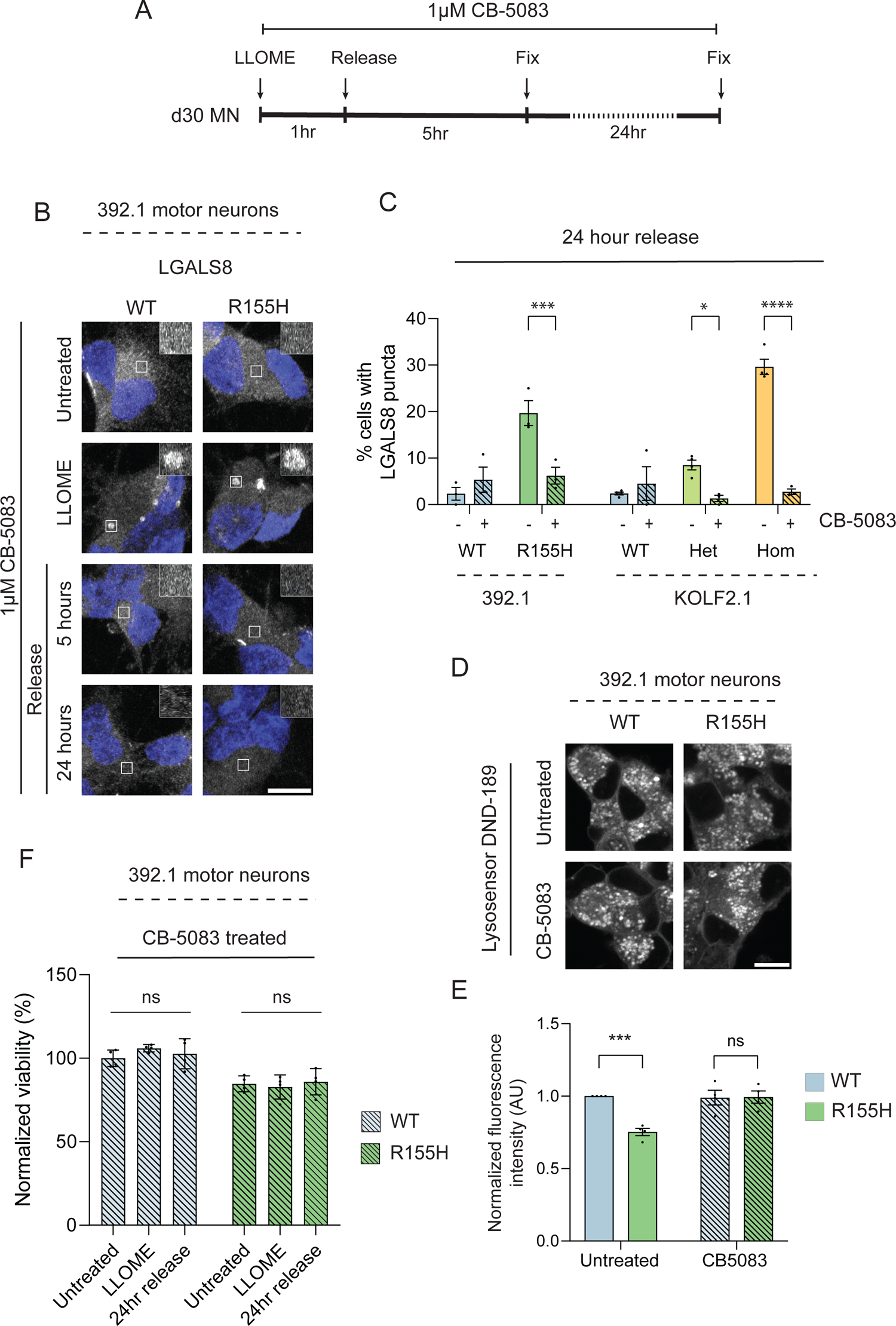
p97 inhibition rescues persistence of damaged lysosomes. A. Schematic of experimental design. B. Representative images LGALS8 puncta in 392.1 motor neurons in untreated, LLOME treated or after release. All conditions contain CB-5083. Scale bar 10µm. C. Quantification of LGALS8 puncta in 392.1 and KOLF2.1 motor neurons 24 hours after LLOME treatment with and without CB-5083 co-treatment. N = 3 independent experiments. Data expressed as means ± SEM. *p<0.05, ***p<0.001, ****p<0.0001; two-way ANOVA with Sidak’s multiple comparison test. D. Representative images of Lysosensor DND-189 live-cell imaging in untreated and CB-5083 treated 392.1 motor neurons. E. Quantification of DND-189 fluorescence in untreated (solid bars) and CB-5083 treated (shaded bars) motor neurons. N = 4 independent experiments. Scale bar 10µm. Data expressed as means ± SEM. ***p<0.001; two-way ANOVA with Sidak’s multiple comparison test. F. Quantification of 392.1 motor neuron viability before, during, and after LLOME and CB-5083 co-treatment. N = 4 independent experiments. Data expressed as means ± SEM. ns-nonsignificant; two-way ANOVA with Dunnett’s multiple comparison test. D30 MN – day 30 motor neurons, LLOME – L-leucyl leucine O-methyl ester.

## Discussion

p97 mutations cause MSP-1 in humans which is a heterogeneous disease involving bone, muscle, spinal cord, and brain. Disease onset and symptom progression is highly variable even between members of the same family. It’s hypothesized that this variability may be due to unknown risk alleles that modulate disease pathogenesis^51^. Due to this heterogeneity, inferring pathways altered due to p97 mutations has been challenging. Previous iPSC-derived motor neuron models of p97 disease have been discordant, with some indicating ER stress and mitochondrial dysfunction^28^ while others attributing disease to aberrant cell cycle protein expression^29^. Similar to previous studies, we found that R155H p97 altered different pathways in different genetic backgrounds. While the KOLF2.1 cell line with R155H p97 displayed mitochondrial deficits (Figures S4B and S4D), the 392.1 patient cell line had no significant alterations to the mitochondrial proteome (Figure S4C).

Unbiased clustering of proteins via WGCNA found that proteins relating to autophagy and endocytosis were altered (Figure 3H). Autophagy has been intensely investigated in ALS as multiple ALS-linked mutations are found in autophagy related proteins^52^. One of these proteins, p62 (SQSTM1), causes multisystem proteinopathy 2 (MSP-2) suggesting that autophagy defects are a common feature in MSPs. It has been well described that mutations in p97 and inhibition of p97 disrupt autophagy in cells^53, 54^ and animals^55, 56^; however, only recently has this disruption has been linked to lysosome repair defects^13, 23^. We identified multiple depleted components of lysosome repair machinery in the few shared DEPs between genetic backgrounds. Interestingly, members of the annexin family, namely ANXA1, ANXA2, and ANXA11 were significantly depleted in mutant motor neurons (Figure 3E). Annexins are calcium-dependent phospholipid binding proteins with diverse functions^57^. A recent study screened annexins for their role in lysosome repair and found that ANXA1 and ANXA2 are uniquely involved in lysosome membrane repair and knockdown of either protein results in delayed repair independent of ESCRT machinery^58^. Mutations in ANXA11 were recently linked to ALS^59^ and a yet unnamed subtype of MSP^60^. ANXA11 contains a low complexity domain which associates with RNA and tethers RNA granules to lysosomes for transport which is thought to be disrupted by disease-linked mutations^61^. While ANXA2 does not have a low complexity domain, it is known to bind mRNA^62^ but whether it plays a similar role to ANXA11 remains to be determined. In addition to annexin proteins, other lipid and/or actin binding proteins were identified as lysophagy DEPs (Figure 3E). Recently, p97 was found to work in conjunction with HSBP1 to extract the actin binding protein CNN2 from damaged lysosomes which is essential for lysophagy^15^. p97 may perform a similar task for additional actin binding proteins; however, further studies are required to determine if these proteins are bona fide p97 substrates,

Galectins are β-galactoside binding proteins that are normally diffusely located in the cytosol but upon LMP bind to exposed luminal lysosomal glycoproteins. While LGALS3 is the best described galectin to localize to damaged lysosomes, LGALS1, LGALS8, and LGALS9 have also been shown to be recruited^44^. Here, we show that motor neurons specifically recruit LGALS8 to damaged lysosomes after LLOME treatment while other galectins including LGALS3 remain diffuse (Figure 4B). Galectins have both shared and unique binding partners that may provide distinct roles for each. LGALS8 has a unique role in regulating mTOR inactivation and has been suggested to promote lysophagy over membrane repair compared to other galectins^63^. Our finding that motor neurons preferentially recruit LGALS8 may indicate that lysophagy dominates membrane repair when faced with LMP; however, further study of lysosome repair and turnover in motor neurons is required to disambiguate the role of LGALS8. Galectin accumulation has been noted in skeletal muscle tissue from patients with p97 mutations^13^ and spinal cord tissue from patients with sporadic ALS^64^. Indeed, changes in galectin levels in cerebrospinal fluid (CSF) and blood have been proposed as potential biomarkers for ALS disease progression^65,66^. Notably, these studies have mainly focused on LGALS1 and LGALS3. Our discovery of LGALS8 as the preferred galectin for lysophagy in spinal motor neurons suggests that LGALS8 should be further investigated as a potential new biomarker.

Lysosome repair is increasingly recognized as an important homeostatic pathway which, when disrupted, may lead to disease^67, 68^. Proper lysosome turnover requires a multi-step process that includes p97-mediated extraction of ubiquitylated substrates^15^. Disruption of this process through inhibition, depletion, or mutation of p97 leads to persistent damaged lysosomes^13,42^; however, to our knowledge, lysophagy has not been studied in human motor neurons. We found that in motor neurons, mutant p97 prevents timely clearance of damaged LGALS8 positive lysosomes (Figure 5). While mutant p97 was recruited to ubiquitylated lysosomes, it did not increase in association with UBXD1 and presumably the ELDR complex (Figures 6 and S6). Without proper ELDR complex assembly, damaged lysosomes are unable to be turned over leading to persistent damaged lysosomes^13^.

In vitro studies of p97 mutations have shown increased ATPase activity leading to the hypothesis that these mutations are gain-of-function^19^. In vivo studies have challenged this notion as knocking out neuronal p97 in mice recapitulates MSP-1 phenotypes^69^. Regardless, p97 inhibitors have been used to attempt to rescue mutant p97-mediated dysfunction with remarkable success^29, 32, 33^. Here, we use CB-5083, a highly specific p97 inhibitor previously used in clinical trials for cancer. We found that treatment with CB-5083 allowed mutant p97 motor neurons to clear LGALS8 puncta at a similar rate as wildtype motor neurons (Figure 7). Interestingly, previous studies have found that p97 inhibition has both decreased LGALS3 puncta clearance^13, 70^ and increased autophagic flux^71^. We found no lysophagy defect in CB-5083 treated wildtype motor neurons (Figure 7B, C). As post-mitotic cells, neurons may be able to better tolerate p97 inhibition than dividing cells such as HeLa or U2OS which have been primarily used in previous studies. In wildtype and mutant neurons, p97 inhibition increased p62 accumulation and in 392.1 neurons, wildtype p62 accumulation surpassed that of mutant motor neurons (Figure S7). The divergence of LGALS8 and p62 puncta here suggests there are additional processes at play that require autophagic clearance. p62 is utilized for ubiquitin-mediated autophagy which includes stress granule clearance and mitophagy: two processes that require p97^72–74^. The additional rescue of lysosomal pH and viability suggests that while p97 inhibition increased p62 accumulation in some conditions, it is not a driver of cell death (Figure 7 and S7).

In conclusion, we establish that LGALS8 is a sensitive marker for lysosomal damage in human motor neurons, and LGALS8 positive lysosomes are not efficiently cleared by mutant p97. The delay in clearance leads to accumulation of p62 and increased cell death. This dysfunction is not due to lack of p97 recruitment but rather decreased association with the ELDR complex component UBXD1. Inhibition of p97 is sufficient to rescue LGALS8 clearance, lysosomal pH and cell survival after lysosomal damage but not p62 accumulation. Our findings add to the increasing evidence that lysosomal homeostasis, particularly lysophagy, are critical components to neuronal health and disruptions in this process lead to disease.

## Supplemental information

**Figure 1S.**
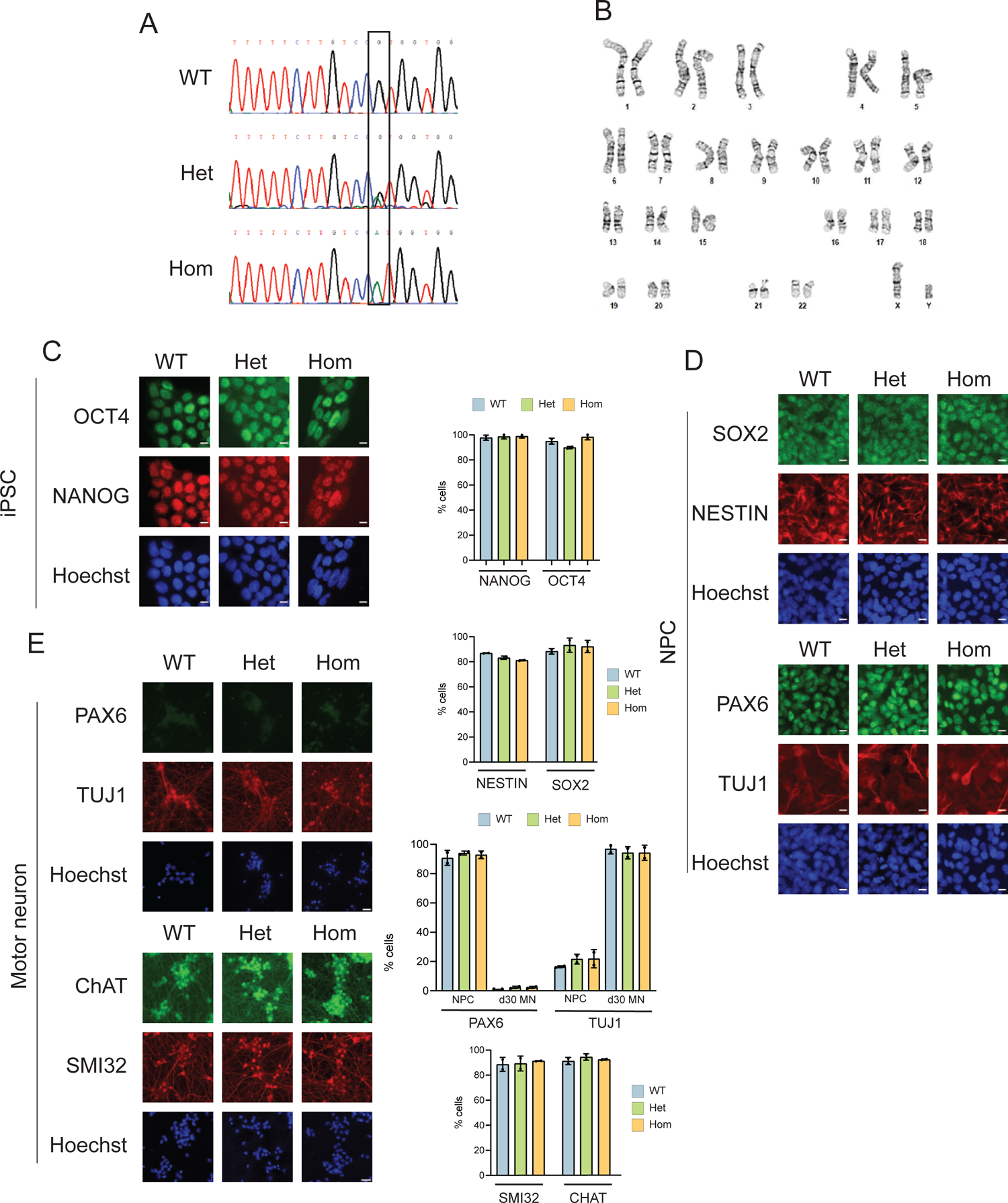
p97 mutations do not affect motor neuron differentiation in the KOLF2.1 cell line. A. Sanger sequencing traces of KOLF2.1 iPSCs confirm WT, heterozygous, and homozygous R155H mutations in the endogenous p97 locus. B. Karyotype of the parental WT KOLF2.1 line shows no macroscopic genetic abnormalities. C. Representative images of iPSCs stained for pluripotency markers OCT4 and NANOG (left panels). Quantification of the percent of cells positive for each marker in each genotype (right). N = 2 independent experiments. Data expressed as means ± SEM. Two-way ANOVA with Dunnett’s multiple comparison test found no significant differences between genotypes. Scale bar 10µm. D. Representative images of NPCs stained for NPC markers SOX2, NESTIN, and PAX6 as well as neuronal marker βIII-tubulin (TUJ1) (right panels). Quantification of the percent of cells positive for each marker (left middle graphs). N = 2 independent experiments. Data expressed as means ± SEM. Two-way ANOVA with Dunnett’s multiple comparison test found no significant differences between genotypes. Scale bar 10µm. E. Representative images of motor neurons stained for the NPC marker PAX6, pan-neuronal marker βIII-tubulin (TUJ1), and motor neuron specific markers choline acetyl transferase (ChAT) and neurofilament heavy chain (SMI32) (left panels). Quantification of the percent of cells positive for each marker (left bottom two graphs). N = 2 independent experiments. Data expressed as means ± SEM. Two-way ANOVA with Dunnett’s multiple comparison test found no significant differences between genotypes. Scale bar 10µm.

**Figure 2S.**
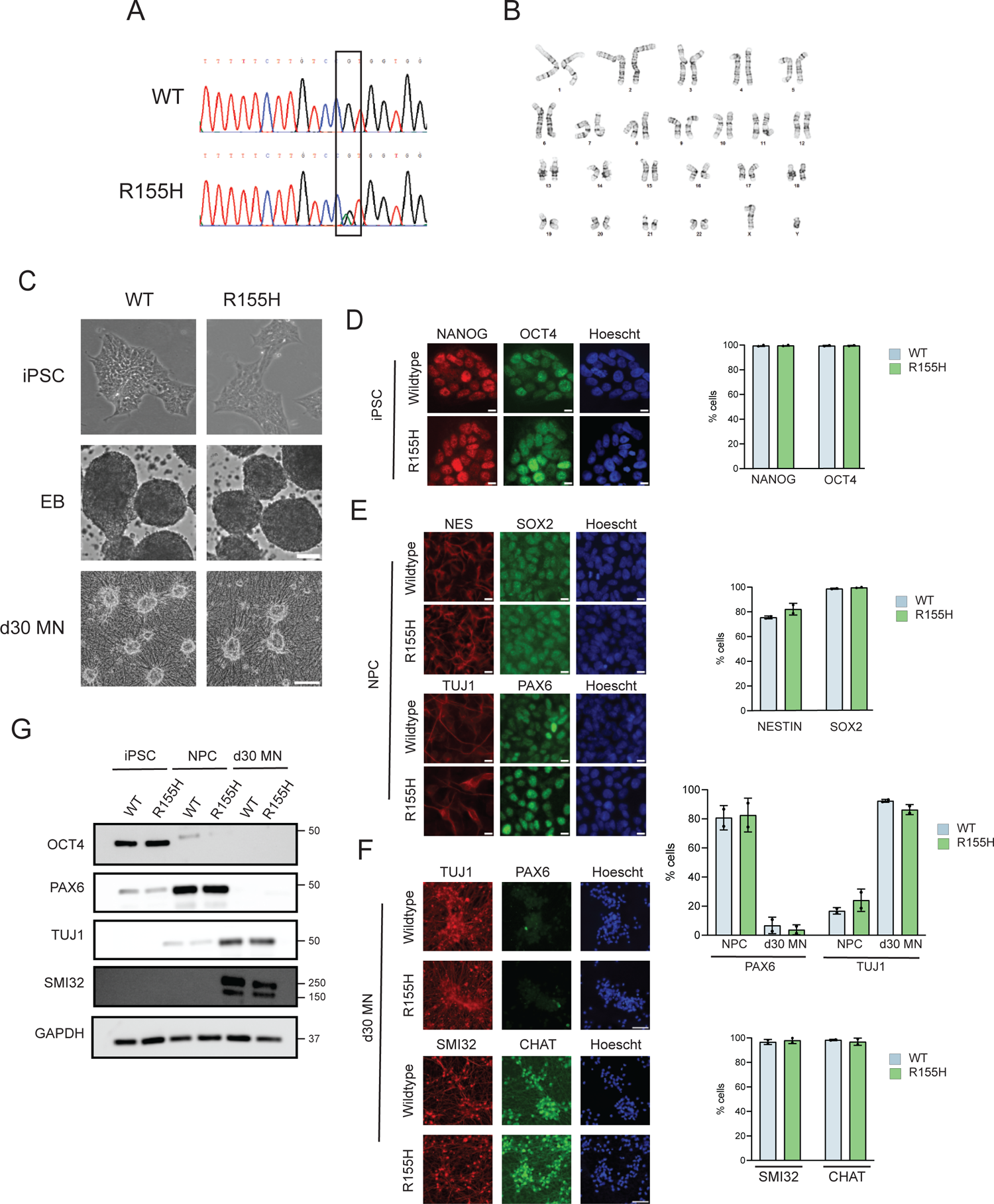
Characterizing of 392.1 and p97 corrected IPSC lines and differentiation. A. Sanger sequencing traces of 392.1 iPSCs confirm WT and heterozygous R155H mutation in the endogenous p97 locus. B. Karyotype of 392.1 iPSCs. C. Phase contrast images of iPSC colonies (top panels), embryoid bodies (middle panels), and motor neurons (bottom panels). Scale bars 200µm (middle) and 20µm (bottom). D. Representative images of iPSCs stained for pluripotency markers OCT4 and NANOG (left panels). Quantification of the percent of cells positive for each marker in each genotype (right). N = 2 independent experiments. Data expressed as means ± SEM. Two-way ANOVA with Sidak’s multiple comparison test found no significant differences between genotypes. Scale bar 10µm. E. Representative images of NPCs stained for NPC markers SOX2, NESTIN, and PAX6 as well as neuronal marker βIII-tubulin (TUJ1) (right panels). Quantification of the percent of cells positive for each marker (left middle graphs). N = 2 independent experiments. Data expressed as means ± SEM. Two-way ANOVA with Sidak’s multiple comparison test found no significant differences between genotypes. Scale bar 10µm. F. Immunoblot of iPSC, NPC, and neuronal markers in each cell type confirms identities. N = 3 independent experiments. G. Representative images of motor neurons stained for the NPC marker PAX6, pan-neuronal marker βIII-tubulin (TUJ1), and motor neuron specific markers choline acetyl transferase (ChAT) and neurofilament heavy chain (SMI32) (left panels). Quantification of the percent of cells positive for each marker (left bottom two graphs). N = 2 independent experiments. Data expressed as means ± SEM. Two-way ANOVA with Sidak’s multiple comparison test found no differences between genotypes. Scale bar 10µm.

**Figure 3S.**
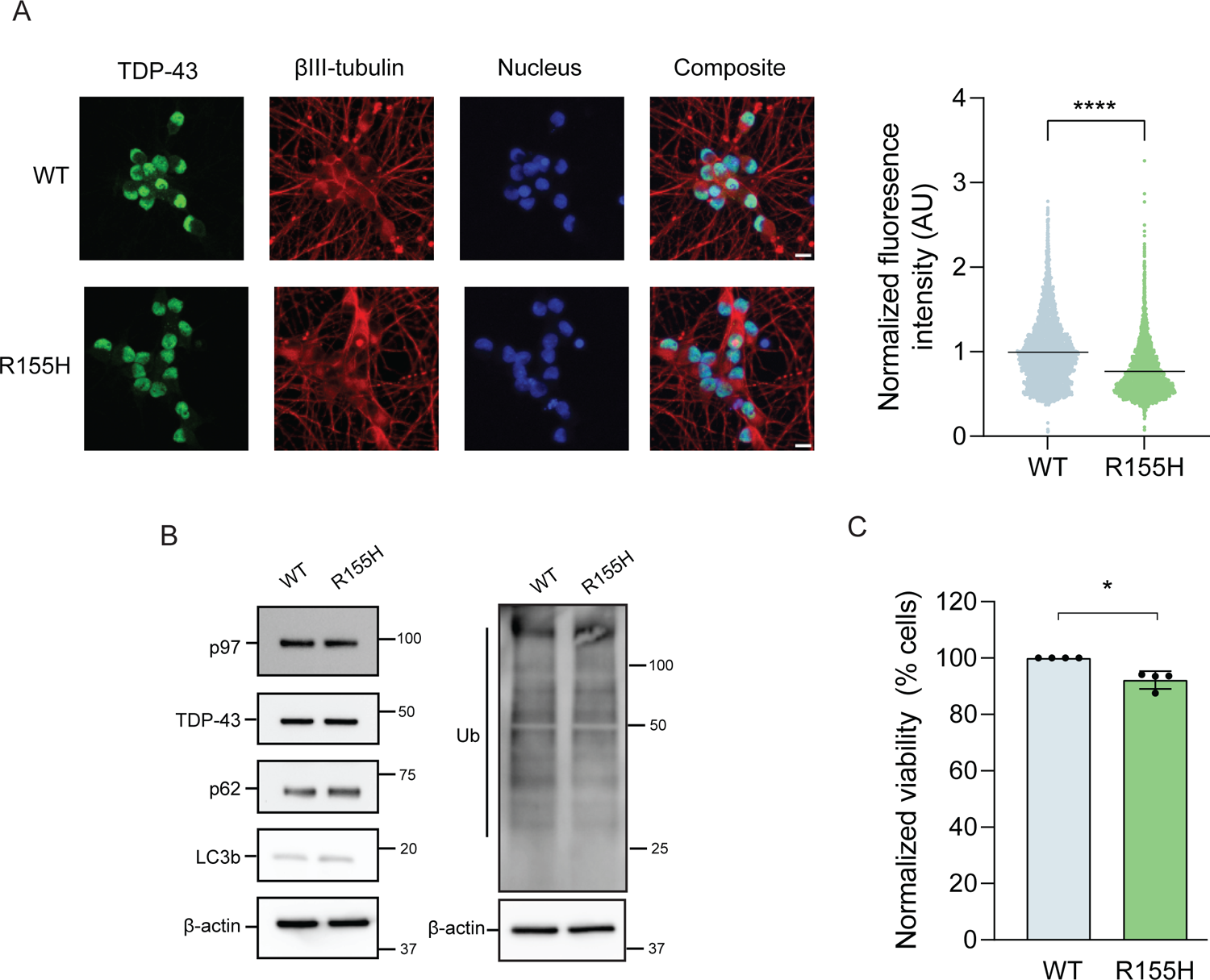
Characterization of ALS phenotypes in 392.1 R155H motor neurons. A. Representative images of TDP-43 immunofluorescence in 392.1 motor neurons (left panels). Quantification of nuclear intensity of TDP-43 staining (right). N = >5000 cells per condition over 4 independent experiments. Data points represent individual cells with a line indicating the mean. ****p<0.0001; unpaired t test. Scale bar 10µm. B. Immunoblots of p97, TDP-43, autophagy markers (p62 and LC3b), and total ubiquitylated proteins (Ub) show no difference between genotypes. N = 3 independent experiments. C. Normalized viability of 392.1 motor neurons following 30 days of differentiation. N = 4 independent experiments. Data expressed as means ± SEM. *p<0.05; unpaired t test.

**Figure 4S.**
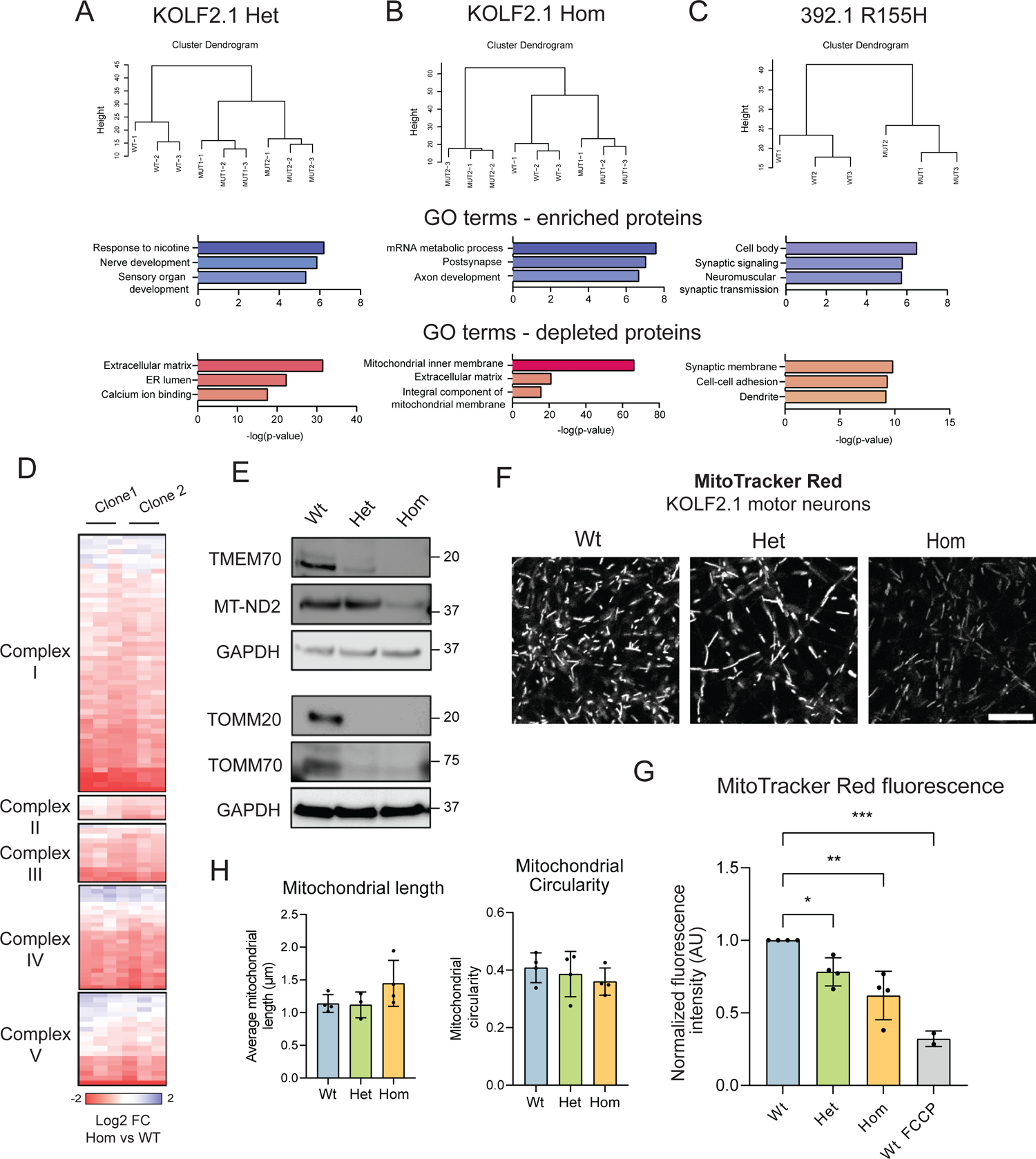
Gene ontology of DEPs in R155H motor neurons reveal genetic background specific differences. A-C. Cluster dendrograms of identified proteins in each experiment indicated (top). Gene ontology terms for enriched and depleted DEPs for each experiment demonstrate divergent proteome changes between cell lines (bottom). D. Heatmap of mitochondrial proteins in KOLF2.1 homozygous motor neurons show wholesale depletion of electron transport chain (ETC) components. E. Validation of the depletion of ETC components TMEM70 and MT-ND2 and translocon components TOMM20 and TOMM70 in KOLF2.1 motor neurons. N = 3 independent experiments. F. Representative images of MitoTracker Red fluorescence in KOLF2.1 motor neurons. Scale bar 10µm. G. Quantification of fluorescence in F. The uncoupling agent carbonyl cyanide p-trifluoromethoxyphenylhydrazone (FCCP) was used to collapse the mitochondrial inner membrane potential as a positive control. N = 4 independent experiments. Data expressed as means ± SEM. *p<0.05, **p<0.01, ***p<0.001; one-way ANOVA with Dunnett’s multiple comparison test. H. Quantification mitochondrial morphology using MitoTracker Red live cell imaging. Data expressed as means ± SEM. One-way ANOVA with Dunnett’s multiple comparison test found no significant differences between genotypes.

**Figure 5S.**
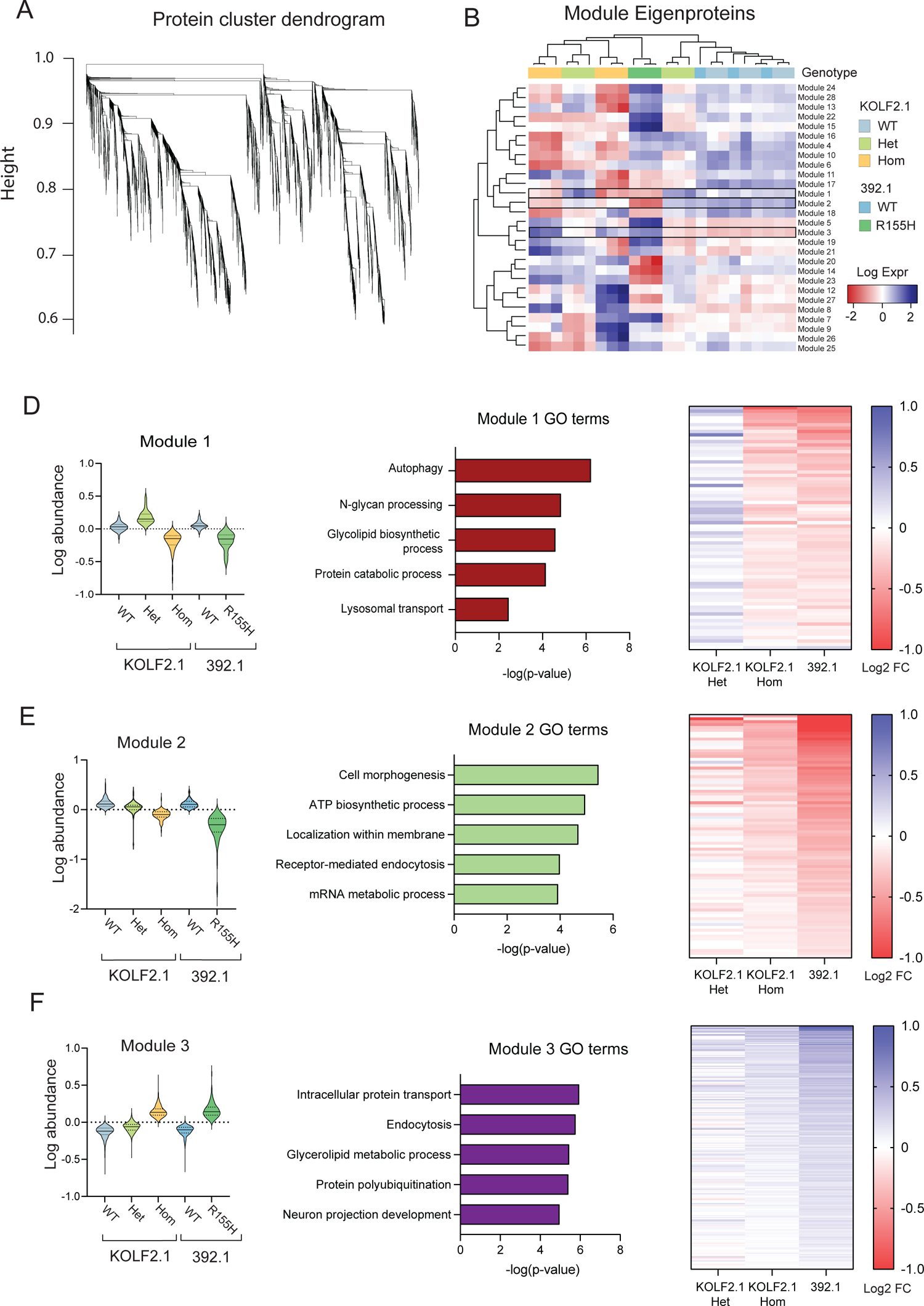
Unbiased clustering analysis of KOLF2.1 and 392.1 motor neuron proteomes identify altered pathways related to autophagy and lysosome homeostasis. A. Weighted gene correlation network analysis (WGCNA) adapted for proteomics experiments was used to cluster all proteins into modules of distinct proteins which are altered in a similar manner between genotypes. B. Heatmap of module eigenproteins for each replicate in each genotype as depicted by color coding above the heatmap. Modules were ranked by concordance and average fold change to identify protein sets changing in a similar manner between cell lines. The top three modules are boxed within the heatmap. D-F. Violin plots of the relative log_2_ abundance of member proteins of the top three modules in motor neurons (left). Top 5 enriched gene ontology terms in each of the modules ranked by significance (middle). Heatmaps of the log_2_ fold change of member proteins in each module for each genotype expressed as mutant vs WT (right).

**Figure 6S.**
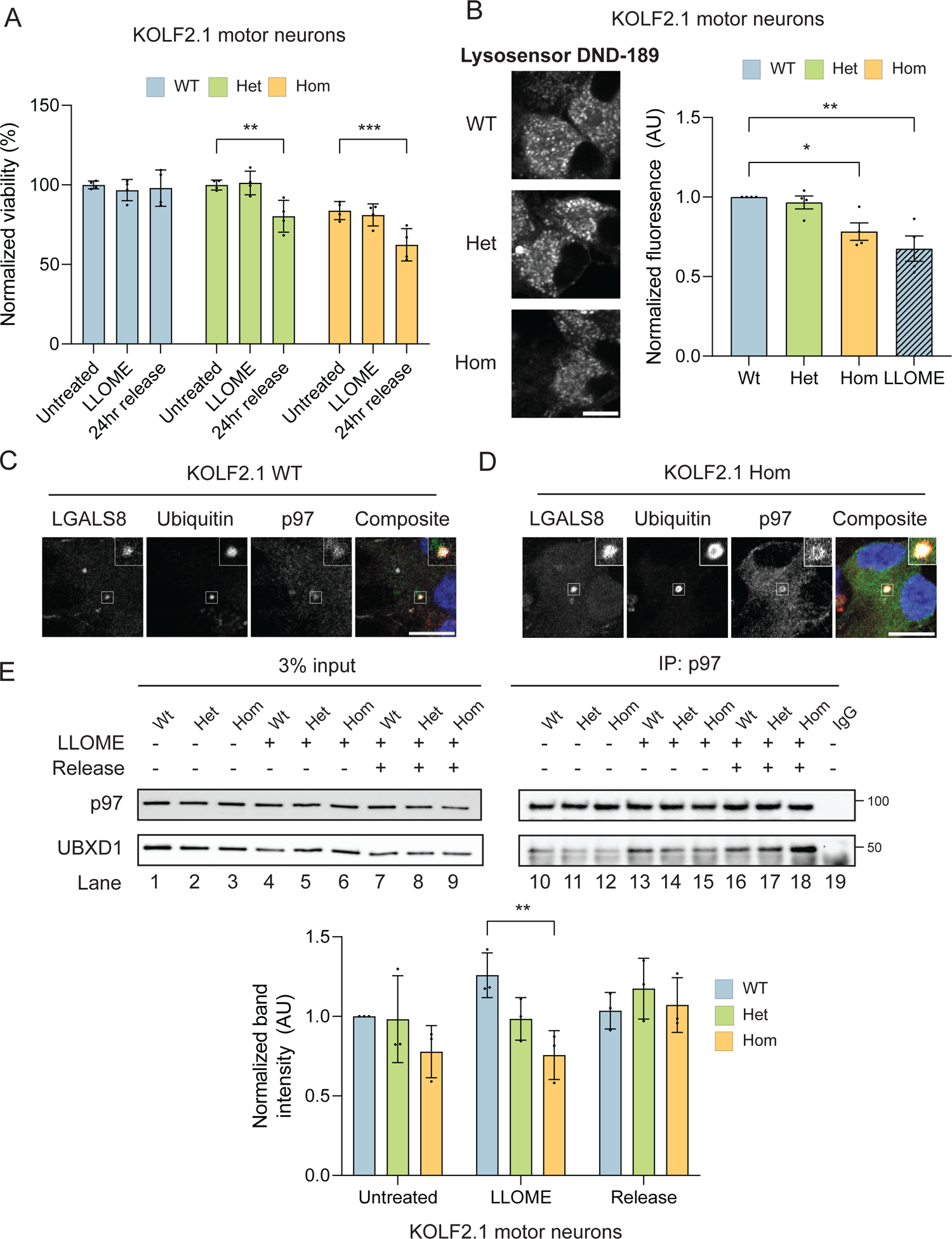
Mutant p97 reduces neuronal resilience to LLOME treatment, increases lysosomal pH, and disrupts association to UBXD1 following lysosomal damage. A. Normalized viability in untreated, LLOME treated or after release in KOLF2.1 motor neurons. N = 4 independent experiments. Data expressed as means ± SEM. **p<0.01, ***p<0.001; two-way ANOVA with Dunnett’s multiple comparison test. B. Representative images of Lysosensor DND-189 live cell imaging in KOLF2.1 motor neurons (left). Quantification of DND-189 fluorescence in untreated and LLOME treated cells. N = 4 independent experiments. Data expressed as means ± SEM. *p<0.05, ****p<0.001; two-way ANOVA with Sidak’s multiple comparison test. Scale bar 5 µm. C-D. Representative images of wildtype (C) and R155H p97 (D) KOLF2.1 motor neurons co-stained with LGALS8, p97, and ubiquitin showing colocalization. N = 3 independent experiments. Scale bars 10µm. E. Immunoblot of endogenous p97 immunoprecipitation in KOLF2.1 motor neurons before, during, and after LLOME treatment (top). Release condition represents 5 hours of recovery. Quantification of UBXD1 band intensities normalized to immunoprecipitated p97 (left). N = 4 independent experiments. Data expressed as means ± SEM. *p<0.05; two-way ANOVA with Dunnett’s multiple comparison test.

**Figure 7S.**
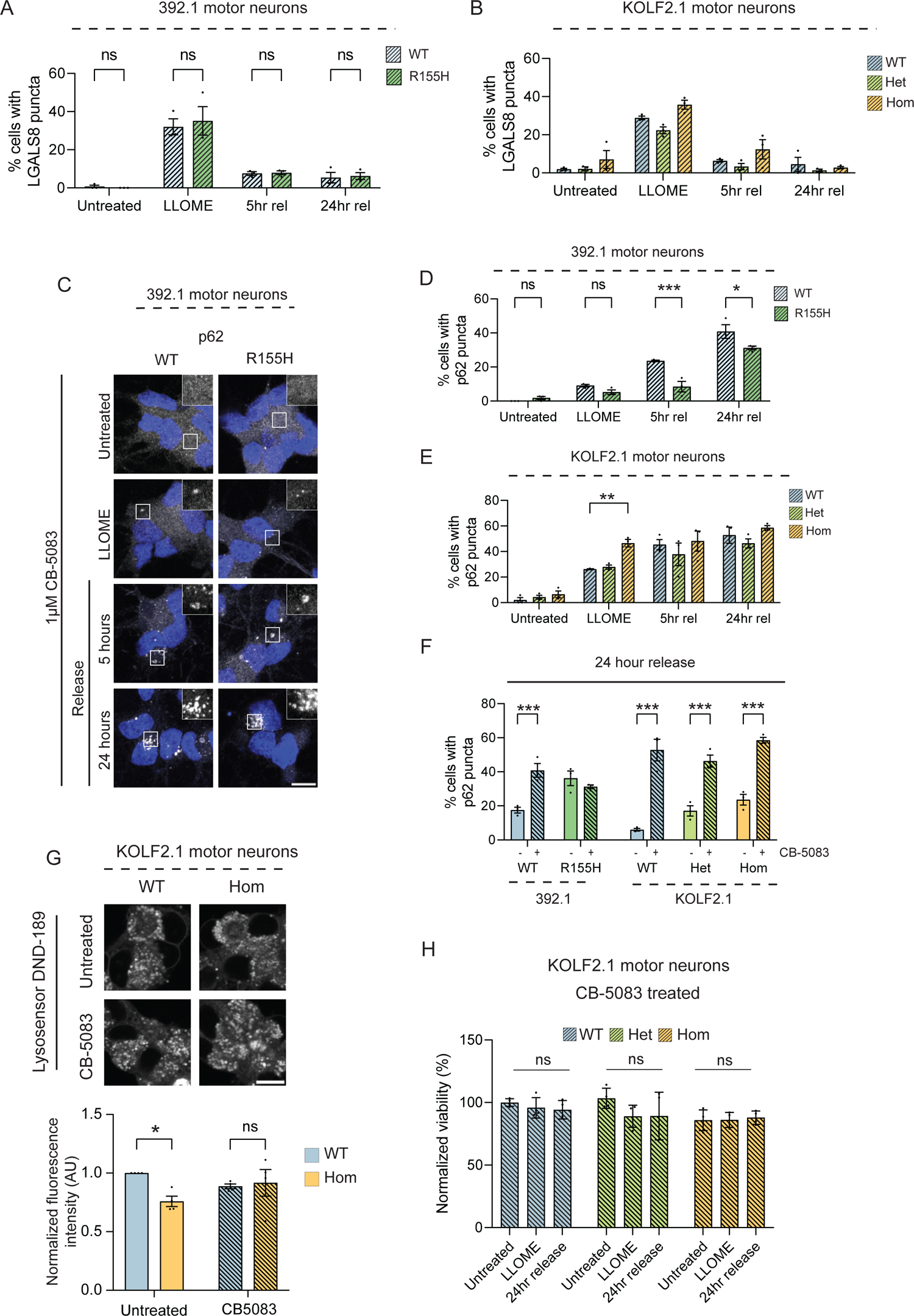
p97 inhibition rescues lysosome defects in motor neurons. A-B. Quantification of LGALS8 puncta in 392.1 (A) and KOLF2.1 (B) motor neurons co-treated with LLOME and CB-5083. N = 3 independent experiments. Data expressed as means ± SEM. ns – nonsignificant. Two-way ANOVA with Sidak’s (A) or Dunnett’s (B) multiple comparison test. C. Representative images p62 puncta in 392.1 motor neurons before, during, and after LLOME treatment. Scale bar 10µm. D-E. Quantification of p62 puncta in 392.1 (D) and KOLF2.1 (E) motor neurons co-treated with LLOME and CB-5083. N = 3 independent experiments. Data expressed as means ± SEM. ns – nonsignificant. *p<0.05, **p<0.01, ***p<0.001; Two-way ANOVA with Sidak’s (D) or Dunnett’s (E) multiple comparison test. F. Comparison of p62 puncta clearance at 24 hours release in motor neurons treated with LLOME (solid bars) or co-treated with LLOME and CB-5083 (shaded bars). N = 3 independent experiments. Data expressed as means ± SEM. ***p<0.001; two-way ANOVA with Sidak’s multiple comparison test. G. Representative images of Lysosensor DND-189 live-cell imaging in KOLF2.1 motor neurons. Quantification of DND-189 fluorescence in untreated (solid bars) and CB-5083 treated (shaded bars) motor neurons. N = 4 independent experiments. Scale bar 10µm. Data expressed as means ± SEM. *p<0.05, ***p<0.001; two-way ANOVA with Sidak’s multiple comparison test. H. Normalized viability of KOLF2.1 motor neurons treated with CB-5083 in untreated, LLOME treated, or after release. Release condition represents 24 hours of recovery. ns – nonsignificant; two-way ANOVA with Dunnett’s multiple comparison test.

## Materials and Methods

### Antibodies and chemicals

Antibodies for LGALS1 (11858-1-AP), LGALS3 (14979-1-AP), PAX6 (12323-1-AP), TDP-43 (10782-2-AP), ATF4 (10835-1-AP), TMEM70 (20388-1-AP), and MT-ND2 (19704-1-AP) were procured from ProteinTech Group. Antibodies for SOX2 (3579T), NANOG (4903T), LC3B (3868T), and Bip (3177T) were procured from Cell Signaling Technologies. Additional antibodies used were TUJ1 (T8660 Sigma), ChAT (ab181023 Abcam), SMI32 (801702 BioLegend), LGALS8 (AF1305 RND Systems), LGALS9 (AF2045 RND Systems), p97 (A300-589A Bethyl (IP) and 612183 BD Biosciences (IF)), ubiquitin (FK2) (04-263 EMD Millipore), LAMP1 (sc-20011 Santa Cruz), GAPDH (sc-47724 Santa Cruz), NESTIN (NBP1-92717SS Novus Biologicals), OCT4 (653701 BioLegend), TOMM20 (sc-17764 Santa Cruz), TOMM70 (sc-390545 Santa Cruz), and p62 (GP62-C Progen Biotechnik). Alex Fluor-conjugated secondary antibodies were from Molecular Probes. Primary antibodies were used at 1:100 for immunofluorescence studies and 1:1000 for immunoblotting. Secondary antibodies were used at 1:5000 for immunofluorescence and immunoblot studies. SB431532 (S1067), Y-27632 (S1049), and SAG1 (S7779) were obtained from Selleck Chem. LDN193189 (SML0559), valproic acid (P4543), BrdU (B9285), PLO (P3655), laminin (L2020), and fibronectin (F2006) were obtained from Sigma-Aldrich. CHIR99021 (4423) was obtained from Tocris. Retinoic acid (sc-200898) was obtained from Santa Cruz. Compound E (565790) was obtained from EMD Millipore. GDNF (450-10), BDNF (450-02), and NT-3 (450-03) were obtained from PeproTech. LLOME (16008) and CB-5083 (19311) were obtained from Cayman Chemicals.

### Induced pluripotent stem cell culture

The KOLF2.1 iPSCs and mutants were acquired from Jackson Laboratories as a part of the iNDI project^30^. 392.1 iPSCs and the CRISPR-corrected line were received from the Weihl laboratory at WUSTL. Patient is a 48 year man with a family history of dementia, weakness and Paget’s disease of the bone. He has history of Paget’s disease of the bone beginning one year prior and presents with subacute onset of slowly progressive right lower extremity and left upper extremity weakness. His EMG/NCS demonstrated active denervation in three body regions including his thoracic paraspinous muscles consistent with a diagnosis of amyotrophic lateral sclerosis. He developed difficulty with breathing at the age of 52 and currently uses non-invasive ventilation at night. He has no symptoms of dementia currently. He has a heterozygous R155H mutation in VCP. iPSCs were maintained in mTeSR (05825 STEMCELL) plus media on Matrigel (354277 Corning) coated plates for no more than 10 passages at 37 °C in humidified incubators. iPSCs were passaged using ReLeSR (05873 STEMCELL) once they reached ∼80% confluency. 10 µM Rho kinase inhibitor Y-27632 (ROCK inhibitor) was added for 24 hours after each split. Cell morphology was routinely monitored for abnormalities. Karyotyping for each cell line was performed by the WiCell Research Institute.

### Neural ectoderm induction and NPC maintenance

We used a modified version of previously published motor neuron protocols^34–36^. Once iPSCs reached 80% confluency, the media was replaced with freshly made induction media (Table 1). Cells were kept in induction media with daily media changes for 3 days. Induction media was replaced with induction-patterning media for an additional 3 days with daily media changes to induce PAX6+ NPCs. At this stage, NPCs were passaged by incubating with 0.05 % trypsin-EDTA (25300054 Thermo) for 4-8 minutes at 37 °C. NPCs were collected using DMEM (SH30243.01 Cytiva) with 10 % FBS (26140079 Gibco) and 1 % P/S (PSL01 Caisson) to inactivate trypsin and centrifuged at 300 x g for 5 minutes. The media was aspirated, and cells were resuspended in fresh expansion media supplemented with 10 µM ROCK inhibitor and plated at 1:4 or 1.5 x 10^5^ cells /cm^2^ on Matrigel coated plates. After 24 hours, the media was replaced with fresh expansion media without ROCK inhibitor. NPCs were banked at this stage in complete expansion media with 10 % DMSO in liquid nitrogen storage. NPCs were thawed, centrifuged at 300 x g for 5 minutes, and plated in expansion media supplemented with 10 µM ROCK inhibitor. 24 hours later the media was replaced with fresh expansion media without ROCK inhibitor. NPCs were passaged for no more than 5 passages or 14 days. Further passaging increases spontaneous differentiation and cultures become more heterogeneous.

**Table 1.**

| Base media | Media type | Component | Concentration |
| --- | --- | --- | --- |
| PIE base | Induction media | SB-431542 | 10uM |
|  |  | LDN-193189 | 1uM |
|  |  | CHIR-99021 | 3uM |
|  | Induction-patterning media | SB-431542 | 10uM |
| | | LDN-193189 | 1 $\mu\text{M}$ |
| | | CHIR-99021 | 3 $\mu\text{M}$ |
| | | Retinoic acid | 2 $\mu\text{M}$ |
| | | SAG1 | 1 $\mu\text{M}$ |
| | Expansion media | CHIR-99021 | 3 $\mu\text{M}$ |
| | | Valproic acid | 5 $\mu\text{M}$ |
| | | Retinoic acid | 0.5 $\mu\text{M}$ |
| | | SAG1 | 1 $\mu\text{M}$ |
| | Patterning media | Retinoic acid | 2 $\mu\text{M}$ |
| | | SAG1 | 1 $\mu\text{M}$ |
| 3M base | Maturation media | Retinoic acid | 1 $\mu\text{M}$ |
| | | SAG1 | 0.5 μM |
|  |  | BDNF | 20 ng/mL |
|  |  | GDNF | 20 ng/mL |
|  |  | NT-3 | 10 ng/mL |
| | | Compound E | 0.2 μM |
| | | Laminin | 1 μg /mL |

**Table 2.**

| Base media | Component | Concentration | Source | Identifier |
| --- | --- | --- | --- | --- |
| Patterning,<br>induction,<br>expansion base<br>(PIE) | DMEM/F12 | 50% | Cytiva | SH30023.FS |
|  | Neurobasal plus | 50% | Thermo | A3582901 |
|  | N-2 supplement | 0.5x | Thermo | 17502048 |
|  | B27 plus supplement | 0.5x | Thermo | A3582801 |
|  | GlutaMax | 1x | Thermo | 35050061 |
|  | NEAA | 1x | Thermo | 11140050 |
|  | Penicillin/streptomycin | 1x | Caisson | PSL01 |
|  | Insulin | 5ug/mL | Sigma | 91077C |
| | β-mercaptoethanol | 100 μM | Gibco | 21985023 |
| Motor neuron<br>maturation<br>media (3M)<br>base | Neurobasal plus | 100% | Thermo | A3582901 |
|  | N-2 supplement | 0.5x | Thermo | 17502048 |
|  | B27 plus supplement | 0.5x | Thermo | A3582801 |
|  | GlutaMax | 1x | Thermo | 35050061 |
|  | NEAA | 1x | Thermo | 11140050 |
|  | Penicillin/streptomycin | 1x | Caisson | PSL01 |
| | Insulin | 5 μg /mL | Sigma | 91077C |
| | β-mercaptoethanol | 100 μM | Gibco | 21985023 |

### Motor neuron differentiation and maturation

NPCs were plated at 1.5 x 10^5^/ cm^2^ in 12-well plates and allowed to grow for 3 days. After 3 days, expansion media was replaced with patterning media for 3 days with daily media changes. After 3 days, the cells were split as described previously, and resuspended in maturation media supplemented with 10 µM ROCK inhibitor. Cells were plated on triple coated PLO/laminin/fibronectin plates at 4 ×10 ^4^/ cm^2^ for biochemistry applications or 1×10 ^4^/ cm^2^ on acid-washed #1.5 glass coverslips (200121 Azer). Half media change was performed the following day with maturation media supplemented with 10 µM ROCK inhibitor. Half the media was replaced 2 days later with maturation media supplemented with 40 mM BrdU to inhibit proliferating cells. Half media changes were performed every 2-3 days for an additional 19 days.

### Western blot

Cells were removed from plates using ReLeSR (IPSCs), 0.05 % trypsin-EDTA (NPCs), or pipetting (motor neurons), centrifuged at 600 x g for 5 minutes, and supernatant removed. Cell pellets were stored at −80 °C until used. Pellets were lysed in RIPA buffer (150 mM NaCl, 50 mM Tris Cl, 0.5 % sodium deoxycholate, 0.1 % SDS, 1% NP-40) supplemented with 1x HALT protease inhibitor (PI-78425 Thermo) and incubated on ice for 10 minutes. Lysates were sonicated at 15% power for 5 seconds, 3 times with 2 second pauses. Lysates were then centrifuged at 21,000 x g at 4 °C for 15 minutes to pellet cell debris. Clarified lysates were transferred to clean microcentrifuge tubes and either frozen at −80°C or used immediately for further assays. Total protein was measured using the BCA assay (23225 Thermo). Protein levels between samples were equalized, mixed with 5x Laemmli buffer (250 mM tris, 25 % ß-mercaptoethanol, 50 % glycerol, 20% SDS, 0.05% bromophenol blue), and boiled for 5 minutes at 95C. An equal amount of sample was loaded and separated on a 12% bis-tris gel and transferred to PVDF membranes (1620177 Bio-Rad) using a Bio-Rad cassette at 70 volts for 2 hours. Membranes were blocked in 5 % milk in tris buffered saline containing 0.1% tween 20 (TBS-T) for 1 hour at room temperature. Primary antibodies were diluted in 2 % BSA (A-420-500 GoldBio) in TBS-T to the indicated concentration and incubated with membranes overnight at 4C with rocking. Membranes were washed 3 times with TBS-T for 10 minutes each while rocking at room temperature. Secondary antibodies were diluted in 2% BSA in TBS-T and incubated with membranes at 1:5000 for 1 hour at room temperature with rocking. Membranes were washed 3 times as described previously, developed with Clarity ECL substrate (1705061 Bio-Rad), and imaged with a ChemiDoc MP Imaging system (BioRad). Blot densities were analyzed using ImageJ.

### Immunoprecipitation

Pellets were lysed in mammalian cell lysis buffer (50 mM Tris Cl, 150 mM NaCl, 0.2 % NP-40, HALT), incubated on ice for 10 minutes, and centrifuged at 21,000 x g for 15 minutes at 4 °C. Clarified lysate was transferred to clean microcentrifuge tubes and total protein was measured using the BCA assay. A small portion of the lysate was set aside as the input, and equal amounts of protein were added to washed protein G beads (20398 Thermo) with the indicated antibody (0.5 µg antibody/ 500 µg of lysate). Samples were incubated at 4 °C overnight while rocking. They were centrifuged at 400 x g for 1 minute at 4C, supernatant was removed, and beads were washed with lysis buffer 3 times. 2X Laemmli buffer was added to samples and boiled for 5 minutes at 95 °C. Samples were centrifuged at 400 x g for 1 minute to pellet the beads and remaining supernatant was analyzed by western blot as previously described.

### qPCR

Total RNA from KOLF2.1 iPSCs, NPCs and d30 motor neurons was harvested and purified using the Quick RNA miniprep kit (R1054 Zymo) following the manufacturer guidelines. Complementary DNA (cDNA) was created for each sample from equals amounts of total RNA using the iScript cDNA synthesis kit (1708890 BioRad). Real-time PCR was performed from cDNA on a StepOnePlus Real-Time PCR System (Applied Biosystems) using the PowerUp SYBR Green Master Mix (A25741 Thermo). Results were quantified using the ΔΔCT method. Signals from samples were normalized against the housekeeping genes, GAPDH or 18s. Primers sequences are as follows:

18s Forward 5’-GGCCCTGTAATTGGAATGAGTC-3’

### 18s Reverse 5’-CCAAGATCCAACTACGAGCTT

Oct4 Forward 5’-GAAACCCACACTGCAGATCA-3’

Oct4 Reverse 5’-CGGTTACAGAACCACACTCG-3’

Pax6 Forward 5’-CCCACACTCTTTATCTCTCACTC-3’

Pax6 Reverse 5’-AGTTGCTGGTGAGAGTTTTCT-3’

ChAT Forward 5’-CCTGCAGTGCATGCGACAC-3’

ChAT Reverse 5’-AAACTGCTGCACAATGGCCT-3’

### Immunofluorescence

Starting on day 9 of differentiation, cells were plated on acid-washed (1 M HCl 4 hours with rocking) # 1.5 glass coverslips that were triple coated (PLO/laminin/fibronectin). Cells were maintained as previously described. Cells were fixed in ice cold 4 % PFA (15710-S Electron Microscopy Sciences) diluted in PBS for 20 minutes. Coverslips were washed once in PBS and incubated in blocking buffer (2 % BSA, 0.2 % Triton-X100 (Figures 1, 2, S1, S2, and S3) or 2 % BSA, 0.4 % saponin (Figures 4, 5, 6, 7, S6, and S7) in PBS) for 1 hour at room temperature. Primary antibodies were diluted to the indicated concentrations in blocking buffer and coverslips were incubated overnight at 4 °C in a humidified chamber. Coverslips were then washed once in blocking buffer and incubated in secondary antibodies diluted to the indicated concentrations in blocking buffer for 1 hour at room temperature. The secondary antibody solution was replaced with Hoechst diluted in PBS and incubated for 5 minutes at room temperature. Coverslips were washed once with PBS and mounted to slides with ProLong Gold antifade mounting media (P36930 Invitrogen).

### LLOME treatment

LLOME was reconstituted to 500mM in sterile DMSO and aliquoted for storage at −20 °C. LLOME was diluted in fresh media immediately before being added to cells. Cells were incubated at the indicated concentrations for the indicated times after which they were washed once with PBS and fixed or fresh media was added for release time points. CB-5083 was reconstituted to 10 mM in sterile DMSO and aliquoted for storage. Fresh aliquots were thawed before each experiment and added directly to complete media to a final concentration of 1µM.

Slides were imaged with either Nikon TI2 Eclipse epifluorescent microscope 40X or 100X objectives using NIS acquisition software (Figures 1, 2, S1, and S2) or Zeiss LSM880 confocal microscope 63X oil objective using ZenBlue software (all others). Image analysis was performed using a modified version of the AggreCount ImageJ macro^75^ or custom ImageJ macros.

### Live-cell imaging

Starting on day 9 of differentiation, progenitors were plated on 4-chambered live-cell imaging dishes (D35C4-20-1-N Cellvis) that were triple coated (PLO/laminin/fibronectin). Cells were maintained as previously described. Cells were treated as previously described. Cells were loaded with LysoSensor DND-189 (L7535 Molecular Probes) or Fluo-4 (F14201 Invitrogen) at 1 µM for 30 minutes or MitoTracker red (M22425 Cell Signaling) at 100 nM for 45 minutes at 37 °C. Cells were then washed once with PBS and the media was replaced with imaging media (complete 3M without phenol red). Cells loaded with Fluo-4 were incubated at room temperature for 20 minutes before imaging. Samples were imaged immediately in a temperature-controlled chamber attached to a Zeiss LSM800 Airyscan confocal microscope. Images were taken at 20 X (Fluo-4) or 63 X (LysoSensor and MitoTracker) magnification. Image analysis was performed in ImageJ using custom macro scripts.

### Electrophysiology

WT and homozygous KOLF2.1 motor neurons were perfused in normal artificial cerebral spinal fluid (nACSF) containing (in mM) 126 NaCl, 26 NaHCO3, 1.25 NaH2PO4, 2.5 KCl, 2 CaCl2, 2 MgCl2, and 10 dextrose (300–310 mosM) and bubbled with 95% O2-5% CO2.

Neurons were recorded under physiological temperature maintained at 33°C (in-line heater; Warner Instruments) and perfused at a high flow rate (∼4 ml/min) throughout the experiment. Input-output curves were generated as previously described^76,77^. Whole cell patch clamp recordings on visually identified motor neurons were performed in the current-clamp configuration using an intracellular recording solution containing (in mM) 130 K-gluconate, 10 KCl, 4 NaCl, 10 HEPES, 0.1 EGTA, 2 Mg-ATP, and 0.3 Na-GTP (pH = 7.25, 280–290 mosM).

The resting membrane potential and input resistance was calculated using Ohm’s law in response to a −100-pA current injection. The presence of spontaneous action potentials was evaluated during a 2-min baseline recording period. The number of action potentials generated in response to a series of 500-ms current injections from 10 to 150 pA in 10-pA steps was measured to generate input-output curves. Series resistance and whole cell capacitance were continually monitored and compensated throughout the course of the experiment. Recordings were eliminated from data analysis if series resistance increased by >20%. For all electrophysiology experiments, data acquisition was carried out using an Axopatch 200B (Axon Instruments) and PowerLab hardware and software (ADInstruments). Data analysis was performed using analysis scripts developed in-house using Python.

### TMT proteomic sample preparation

Samples were collected by aspirating media and pipetting cells off with ice cold PBS. Cells were spun at 600 x g for 5 minutes at 4 °C, the supernatant was removed, and cells were stored at −80 °C. Cell pellets were lysed with 8 M urea supplemented with 1x HALT in 200 mM EPPS (E0276 Sigma) by pipetting up and down 10 times. Samples were passed through a 26g syringe 10 times to shear membranes and centrifuged at 21,000 x g for 10 minutes to pellet debris. The supernatant was transferred to clean microcentrifuge tubes and total protein was estimated using the BCA assay. 100 µg of protein was aliquoted from each sample and the volumes were made equal with lysis buffer. Samples were reduced with 5 mM TCEP for 30 minutes, alkylated with 14 mM iodoacetamide for 30 minutes, and the reaction was quenched with 5 mM DTT for 15 minutes all at room temperature in the dark. Protein was precipitated by adding 400 µL methanol, 100 µL chloroform, and 300 µL water and centrifuging 21,000 x g for 2 minutes at room temperature. The organic and aqueous layers were aspirated, and the remaining protein was allowed to air dry for 5-10 minutes. The protein precipitate was resuspended in 100 µL of 200 mM EPPS. 1 µg/µg of Lys-C (129-02541 Wako) was added to each sample and incubated overnight at room temperature with shaking. Samples were further digested with 1 µg/µg trypsin (90305 Thermo) per sample for 6 hours at 37 °C with shaking. 30 µL of anhydrous acetonitrile was added to each sample. Samples were labeled with 10-plex TMT labeling reagents (90110 Thermo) at 1:10 for 1 hour at room temperature. The labeling reaction was quenched with 10uL 5% hydroxylamine in 200mM EPPS for 15 minutes. TMT-labeled peptides from each sample were combined in equal amounts. The pooled sample was dried under vacuum.

The sample was resuspended in 5% formic acid for 15 minutes. The peptide mix was desalted using C18 solid-phase extraction (SPE) (WAT036945 Waters). This sample was then fractionated using off-line basic pH reverse-phase fractionation via high performance liquid chromatography (HPLC). The peptide mix was fractionated in 96 fractions which were combined into a total of 24 fractions. These fractions were further desalted using STAGE tips made from C18 resin and p200 pipette tips and died under vacuum. They were reconstituted in 5% acetonitrile / 1% formic acid for LC-MS/MS processing. An Orbitrap Lumos mass spectrometer coupled with a Proxeon NanoLC-1000 UHPLC was used for data collection. Proteins were identified using all entries from the Human UniProt Database (2018). Peptide-spectrum matches were adjusted to a 1% false discovery rate. Raw protein abundances were determined by summing reporter ion counts across all matching peptides. The abundances were adjusted for protein loading and scaled to generate a relative abundance measurement.

### Proteomic data analysis

Log_2_ fold changes and adjusted p values were calculated using a modified linear modeling for microarray data as an empirical Bayes procedure using a custom R script^78^. Weighted gene co-expression analysis (WGCNA) was performed as previously described using a modified version of the WGCNA R script^41^. In brief, relative abundances were normalized using sample loading normalization and internal reference scaling to eliminate variation between proteomics experiments. Normalized abundances were used to build a topological overlap distance matrix which was clustered using the TOM-based dissimilarity method. Modules were identified via a dynamic tree cutting algorithm. These modules were ranked based via concordance between cell line (KOLF2.1 Hom and 392.1 R155H) and average log_2_ FC. The three highest ranked modules were used for gene ontology analysis using Metascape to identify enriched pathways^79^.

### Statistics and reproducibility

For all experiments, n ≥ 3 or more biological replicates for each condition examined except immunofluorescence characterization experiments in figures 1, S1, and S2 (n = 2). Fold changes, SEM, SD, and statistical analyses were performed using GraphPad Prism version 9.4.1 for Windows (GraphPad Software). Statistical tests and n values are mentioned in the figure legends.

## Acknowledgements

We thank members of the Raman lab for critical reading of the manuscript. This work is supported by the NIH grants GM127557 and NS123631 to M.R. M.A.J is an IRACDA scholar funded by GM133314. This work was funded in part by NIH grant GM67945 (S.P.G.) and GM132129 (J.A.P.), AA026256, NS105628, NS102937, MH128235, and MH122379 (J.M) and AG031867 and AR073317 (C.C.W).

## Respective Contributions

J.K and M.R conceived the studies. J.K performed all studies and data analysis, M.J performed qRT-PCR and assisted in validation studies. C.P and J.M performed electrophysiology and analysis, J.P and S.P.G assisted with proteomic studies. C.C.W generated the patient iPSC line. J.K and M.R wrote the manuscript.

## Request for reagents

Please contact the corresponding author, M.R for reagent requests.

## Data availability

All raw proteomic data will be made available on public servers upon acceptance of the manuscript. Any other data is available from the corresponding author upon request.

